# Direct Effects of Cigarette Smoke in Pulmonary Arterial Cells alter Vascular Tone through Arterial Remodeling and Kv7.4 Channel Dysregulation

**DOI:** 10.1101/555953

**Authors:** J Sevilla-Montero, D Labrousse-Arias, C Fernández-Pérez, B Barreira, G Mondejar-Parreño, A Cogolludo, MJ Calzada

## Abstract

Chronic obstructive pulmonary disease (COPD) is a widespread disease, with no curative therapies nowadays. Exposure to cigarette smoking is considered the chief leading cause of COPD. Current drugs therapies improve patient quality of life, however they do not revert the progression of the disease. Therefore, a deeper study of the cellular and molecular mechanisms that underlie this pathology is required to be able to carry out targeted and effective treatments. Although the effects of cigarette smoke in the progressive deterioration of the airway have been extensively studied in COPD patients, its effects on pulmonary vasculature have been unexplored, due to the classic conception that vascular damage is a consequence of alveolar hypoxia and loss of capillary bed. In this paper, we aimed to study the effects of cigarette smoke extract (CSE) in regulating pulmonary arterial cells phenotypic modulation, in particular the effects in fibroblasts (hPAFib) and smooth muscle cells (hPASMC), and in murine pulmonary arteries. Our results demonstrated that CSE exposure had direct effects on hPAFib and hPASMC, promoting a senescent phenotype that in turn contributed, through the secretion of inflammatory molecules, to increase the proliferative potential of non-exposed cells. CSE also increased total ROS levels in hPAFib and hPASMC, and upregulated NADPH oxidase subunits NOX1 and p22^phox^. Most importantly, CSE affected cell contractility and dysregulated the expression and activity of voltage-gated K^+^ channel Kv7.4. This contributed to limit vascular responses impairing vasoconstriction and endothelium-dependent and independent relaxation.

## Introduction

Chronic Obstructive Pulmonary Disease (COPD) is currently the fourth leading cause of death in the world [1, 2], but is expected to be the third leading cause of death by 2020. The advanced stage of the disease is characterized by a progressive development of non-reversible airflow limitation, loss of functionality, and a great number of exacerbations, all these leading to a significant deterioration in the patient quality of life. Environmental exposure, primarily to cigarette smoking, is considered the chief leading cause of COPD. The use of bronchodilator and corticoid drugs improve the patient quality of life by reducing dyspnea and increasing exercise tolerance, however they do not revert the progression of the disease and current treatment remains unsatisfactory. Since there are no therapies to halt its progression it is important to understand the pathophysiological, histological, cellular and molecular mechanisms that underlie this pathology, in order to be able to carry out targeted and effective treatment and prevention.

Prominent forms of the disease include chronic bronchitis and emphysema but some patients also develop pulmonary hypertension (PH) which is the end result of interaction between pulmonary vascular tone and a complex series of cellular and molecular events termed vascular remodeling [3, 4]. The classic conception propose that this vascular remodeling is a consequence of alveolar hypoxia and loss of capillary bed. This induces a continuous vasoconstriction and a state of hypoxemia with a concomitant vascular remodeling, triggering PH. However, several pieces of evidence pointed to vascular damage as an independent process that may be previous to alveolar damage. In this respect, there are patients who develop PH prior to hypoxemia, and endothelial dysfunction with mild COPD [5-7]. This has also been demonstrated in the pulmonary vasculature of necropsies from healthy smokers and patients with COPD without hypoxemia and in animal models chronically exposed to tobacco smoke [8-11]. These findings suggest that vascular remodeling may be driven by mechanisms affecting the pulmonary vasculature independent of hypoxemia. Additionally, they point out that cigarette smoke plays an important role in pulmonary vascular remodeling. In this respect, results from animal models and human cells have shown that cigarette smoke induces proliferation in rat pulmonary artery smooth muscle cells, fibroblasts and endothelial cells and in human airway and aortic smooth muscle cells [12-14]. Conversely, other studies demonstrate that exposure to cigarette smoke inhibits proliferation of lung fibroblasts [15] and human pulmonary artery endothelial cells (hPAEC) [16], while inducing their senescence, which leads to improved smooth muscle cell (hPASMC) proliferation and migration [17]. Interestingly, the existence of a combined phenotype of both proliferative and senescent cells in pulmonary arteries from COPD patients has also been shown [18]. Although cellular senescence in lung cells from smokers and COPD patients has been attributed to the induction of p16 [19, 20], other authors have shown that deletion of p16 is not sufficient to mediate protection against senescence, suggesting that additional mechanisms may be involved [21]. Additionally, ROS production, known to induce senescence, is increased in COPD fibroblasts [22]. However, considering the variability of results with respect to cellular behavior in the arteries of patients with COPD, it is necessary to identify the effector agents of these processes as well as the underlying cellular and molecular mechanisms.

K^+^ channels present in pulmonary artery smooth muscle cells (PASMCs) make a major contribution in the regulation of pulmonary vascular smooth muscle tone. Their activation turns the membrane potential more negative, and as a consequence, may contribute to vasodilation by causing voltage-gated calcium channels closure. In fact, activation of several K+ channels, including large conductance calcium-dependent potassium channels (BKCa) or voltage-gated K channels (Kv; especially Kv1 and Kv7 channels) has been suggested to contribute to the pulmonary vasodilation effects induced by the nitric oxide (NO) pathway [23, 24] (Mondejar-Parreño et al., 2019). In contrast, reduced K+ channel performance induces a more depolarized membrane potential in PASMCs, which leads to increased intracellular calcium, vasoconstriction, hypertrophy and proliferation [25-29]. Thus, impairment of K+ channels is considered a hallmark in the pulmonary vascular alterations associated with pulmonary hypertension and other respiratory conditions such as asthma and COPD [30-35]. However, the possible alteration of K+ channels by CSE in PASMC has been essentially unexplored.

The effects of cigarette smoke in the progressive deterioration of the airway have been extensively studied in COPD patients. However, the effects on pulmonary vasculature have been neglected, due to the classic conception that vascular damage is a consequence of alveolar hypoxia and loss of capillary bed. In this paper, we aimed to study the effects of cigarette smoke extract (CSE) in regulating pulmonary arterial cells phenotypic modulation and in particular the effects in hPAFib and hPASMC. Our results demonstrated that CSE exposure had direct effects on hPAFib and hPASMC, promoting a senescent phenotype that in turn contributed, through the secretion of inflammatory molecules, to increase the proliferative potential of non-exposed cells. In addition, CSE affected cell contractility and dysregulated the expression and activity of voltage-gated K^+^ channel Kv7.4. This ultimately affected vascular responses impairing vasoconstriction and endothelium-dependent and independent relaxation.

## METHODS

### Cell culture

Primary human pulmonary artery adventitial fibroblasts (hPAFib) and smooth muscle cells (hPASMC) were obtained from ScienCell Research Laboratories (#3120 and #3110, respectively). Cells were cultured, following manufacturer’s recommended specifications, in fibroblast or SMC medium (ScienCell, #2301 and #1101) containing 2% fetal bovine serum, 1X fibroblast or SMC growth supplement (ScienCell, #2352 and #1152), 100 U/mL penicillin and 100 μg/mL streptomycin. Cells were maintained at 37 °C in a humidified atmosphere of 5% CO_2_. To avoid the exposure to volatile substances from the smoke, untreated and CSE-challenged cells were always kept in different incubators.

### Preparation of cigarette smoke extract (CSE)

CSE was prepared from commercial Marlboro Red cigarettes (Philip Morris Brand Sàrl Neuchâtel, Switzerland). Briefly, the smoke from the complete consumption of five cigarettes was continuously bubbled into 50 mL of culture medium using VACUSAFE aspiration system (INTEGRA Biosciences AG) at a vacuum pump flow rate of 8 L/min. The CSE solution was filtered through a 0.22 μm-pore system, immediately aliquoted and kept at −20 °C until used. To ensure a similar preparation amongst different batches, CSE concentration was measured spectrophotometrically at 320 nm wavelength, showing a mean optical density of 0.114 ± 0.006. This solution was considered to be 100% CSE and diluted to obtain the desired concentrations for each experiment.

### CFSE proliferation assay

Cells were stained with 0.5 μM of the fluorescent probe carboxyfluorescein succinimidyl ester, CFSE (CellTrace™, Thermo Fisher Scientific, C34554) in Hank’s Balanced Salt Solution (HBSS) (Lonza, BE10-527F) for 20 minutes at 37 °C. Afterwards, complete medium was added for another 5 minutes to stop the staining process and cells were allowed to attach on 24-well plates before CSE challenge. Cells were trypsinized and median CFSE fluorescence intensity was measured on a FACSCanto™ II cytometer (BD Biosciences) illuminating with Ar 488 nm laser, right before CSE exposure (day 0) or 72 hours later. Results were expressed as the CFSE fluorescence lost with respect to day 0 condition and normalized to untreated cells fluorescence levels. For conditioned medium experiments, cells were CSE-exposed for 48 h, changed to fresh culture medium and the secretome was collected during the following 48 hours. This conditioned medium was then added to CSE-unexposed hPAFib and hPASMC and proliferative response was assessed 72 hours later.

### X-Gal staining

For β-galactosidase activity detection, Senescence Cells Histochemical Staining Kit (Sigma Aldrich, CS0030) was used according to manufacturer’s instructions. Briefly, after CSE exposure for 48 hours, cells were fixed with 2% formaldehyde – 0.2% glutaraldehyde buffer for 10 minutes at room temperature and incubated in X-Gal staining solution at 37 °C overnight. The development of blue color, which indicates β-galactosidase activity, was analyzed in phase contrast images of three random fields per condition with a Nikon Eclipse TS100 inverted microscope using a Nikon air objective CFI Achro ADL10X/0.25 and NIS-Elements software. Results were expressed as percentage of β-galactosidase^+^ cells out of total cells for each CSE dilution.

### Measurement of cell surface area

For hypertrophic phenotype determination, hPAFib and hPASMC were challenged with different CSE dilutions for 24 hours and phase contrast imaged afterwards. Total apparent surface from each cell was quantified using ImageJ 1.51 software (National Institutes of Health, Bethesda, MA, USA).

### Protein expression by western-blot analysis

Lysates of hPAFib and hPASMC grown to 90% confluence in 35 mm plates were prepared in reducing 2X Laemmli buffer, boiled at 95 °C for 10 minutes, electrophoretically separated by sodium dodecyl sulphate polyacrylamide gel electrophoresis and transferred onto 0.45 μm nitrocellulose membranes (GE Healthcare Life Sciences, 10600003). Total protein bands were reversibly stained with Fast Green FCF (Sigma-Aldrich, F7252) and visualized on ImageQuant LAS 4000 (GE Healthcare Life Sciences) for a later quantification of total protein. Blots were washed and then incubated with 5% nonfat dry milk in Tris buffered saline–0.1% Tween 20 for 30 minutes at room temperature. Afterwards blots were probed overnight at 4°C with the following primary antibodies: anti-p21 clone SXM30 (BD Pharmingen, 65961a), anti-p53 clone DO-1 (Santa Cruz, sc-126), anti-p16 clone EPR1473 (Abcam, ab108349), polyclonal anti-Kv1.5 (Alomone Labs, APC-004), polyclonal anti-Kv7.4 (Alomone Labs, APC-164), monoclonal anti-Kv7.5 (Santa Cruz, sc-293305) and polyclonal anti-BKCa (Alomone Labs, APC-151). Horseradish peroxidase-conjugated anti-mouse (Dako, P0260) or anti-rabbit (GE Healthcare Life Sciences, NA934V/AG) antibodies were added for 1 hour at room temperature and protein signal was then visualized using Immobilon Forte (Millipore, WBLUF0500) on ImageQuant LAS 4000 (GE Healthcare Life Sciences).

### Real-time PCR analysis

Cells were grown to 90% confluence in 100 mm plates and total RNA was isolated using TRI Reagent (Molecular Research Center Inc., TR118). RNA (1 μg) was reverse transcribed to cDNA with MultiScribe™ Reverse Transcriptase (Applied Biosystems, 4308228) in a final volume of 20 μl. cDNA (1 μl) was amplified with specific primer pairs, and quantitative PCR was performed using Power SYBR green PCR Master Mix (Promega, A6001) and the specific primer pairs shown in Supplementary table S1. Data were analyzed by the 2^−ΔΔCt^ method using StepOne Plus Software (Applied Biosystems) and all values were controlled with *ACTB* gene expression levels.

### Propidium iodide cell cycle assay

After 24 hours of CSE treatment, cells were trypsinized, fixed with ice cold 66% ethanol in PBS and stored until use at 4 °C. On the day of analysis, the cells were centrifuged, resuspended in propidium iodide (Immunostep, ANXVDY-100T) and 500 U/mL RNase A (Sigma-Aldrich, R6513) solution and incubated at 37 °C for 30 minutes. Following incubation, cells were kept on ice and immediately acquired on a FACSCanto™ II cytometer (BD Biosciences) illuminating with Ar 488 nm laser. Interval gates were placed on the detected peaks of a propidium iodide histogram plot and percentages from each gate representing G0/G1, S and G2/M phases were estimated with FlowJo 8.7.3 software (Tree Star Inc., Ashland, OR, USA).

### Reactive oxygen species quantification

ROS were quantified with the fluorescent probe dihydroethidium (DHE) (ThermoFisher Scientific, D1168). Following 24-hour treatment with CSE, cells were incubated with 10 μM DHE probe in HBSS for 20 minutes at 37 °C. Afterwards, cells were trypsinized and median DHE fluorescence was measured on a FACSCanto™ II cytometer (BD Biosciences) illuminating with Ar 488 nm laser. Specific changes in DHE signal intensity were quantified by subtracting fluorescence from PAFib and PASMC treated with DMSO as vehicle control, and normalized to non CSE-exposed cells.

### Collagen gel contraction assay

To evaluate cell contraction, hPASMC were mixed with 1 mg/mL collagen type I from bovine skin (Sigma Aldrich, C4243) and 4 mM NaOH in their respective media and placed in 24-well plates in a volume of 500 μl containing 1.2·10^5^ cells/well. Mixture was allowed to gel at 37 °C for at least 30 minutes and carefully detached from the well wall using the end of a tip before adding 500 μL of medium on top. Next day, cells were challenged with CSE and gels were imaged at 24, 48 and 72 hours afterwards on ImageQuant LAS 4000 (GE Healthcare Life Sciences). Images were analyzed by measuring the gel surface area using ImageJ 1.51 software. Data were presented as percentage of initial gel area along time.

### Traction force microscopy (TFM) imaging

The TFM experiments were carried out according to the protocol previously described by Plotnikov SV and collaborators [36]. Briefly, Polyacrylamide gels with a Young’s modulus of 5 kPa, and a Poisson ratio of 0.5 were prepared with 2.5% (v / v) of 0.2 μm crimson-labelled fluorescent microspheres FluoSpheresTM (ThermoFisher Scientific F8806) embedded in their interior. Gels were then functionalized overnight at 4°C with 250 μg/mL fibronectin using the photoactivatable crosslinker Sulfo-SANPAH (Thermo Fisher Scientific, 22589). After 24 hours CSE exposure, cells were stained for 20 minutes with 5 μM CellTrace™ CFSE and immediately placed on top of functionalized polyacrylamide gels. When the cells attached, images of the gels were obtained in a Leica SP5 confocal microscope using a Leica oil immersion objective HC PL APO 40X/0.75 CS2, illuminating with Ar 488 nm and HeNe 633 nm lasers. Cells were continuously recorded for 10 minutes while full gel relaxation was accomplished by a treatment with 0.05% trypsin-EDTA (Thermo Fisher Scientific, 25300-062).

### Beads displacement and traction stress reconstruction

Confocal images were collected using Leica TCS SP5 software (Leica Microsystems, Mannheim, Germany). Subsequent analysis was performed on ImageJ 1.51, following a previously described computational strategy [37]. Iterative particle image velocimetry and regularized Fourier transform traction cytometry methods were implemented to calculate fluorescent beads displacement and traction stresses exerted by the cells [38]. Critical computational parameters such as interrogation window size, correlation threshold and FTTC regularization factor were set equal to 16 pixels, 0.6 and 8·10^−10^, respectively. MATLAB R2015b software was finally used to calculate cell surface and total traction exerted and to assign a color code to the stress fields.

### Vascular contractility measurement

Pulmonary arteries (PAs) from male C57BL/6 WT mice (Jackson Laboratory, stock number 000664) were carefully dissected free of surrounding tissue, cut into rings (1.8-2 mm length) and overnight-exposed to CSE. Afterwards, vessel segments were mounted on a wire myograph in the presence of Krebs physiological solution. Contractility was recorded by an isometric force transducer and a displacement device coupled with a digitalization and data acquisition system (PowerLab, Paris, France). Preparations were ?rstly stimulated by raising the K^+^ concentration of the buffer (to 80 mM) in exchange for Na^+^. Vessels were washed three times and allowed to recover before a new stimulation. Arteries were then stimulated with 10^-5^ M serotonin (5-HT, Sigma) and treated with increasing concentrations of the endothelium-dependent vasodilator acetylcholine (ACh, from 10^−9^ M to 10^−5^ M, Sigma), the nitric oxide donor sodium nitroprusside (SNP, from 10^−11^ M to 10^−5^ M, Sigma) or the Kv7 channel activator retigabine (10^−8^ M to 3·10^−5^ M, Sigma). Some PAs were treated with XE991 (3·10^-8^-3·10^-6,^ Sigma) before the stimulation with 5-HT and then the relaxation induced by SNP was tested.

### Patch clamp

Following overnight-exposure to CSE, SMC were isolated by enzymatic digestion from murine PAs rings as previously described [39]. Membrane currents were recorded at room temperature with an Axopatch 200B and a Digidata 1322A (Axon Instruments, Burlingame, CA, USA) using the whole-cell configuration of the patch-clamp technique. Cells were superfused with an external Ca^2+^-free HEPES solution (120 mM NaCl, 5 mM KCl, 5 mM Na_2_ATP, 10 mM MgCl_2_, 5.6 mM glucose, and 10 mM HEPES) and a Ca^2+^-free pipette (internal) solution containing: 130 mM KCl, 1.2 mM MgCl_2_, 5 mM Na_2_ATP, 10 mM HEPES, 10 mM EGTA. Kv currents were evoked following the application of 200 ms depolarizing pulses from −60 mV to +60 mV in 10 mV increments. Cell capacitance was calculated from the integral of the capacitive transient current elicited by 10-mV hyperpolarizing pulses from a holding potential of −70 mV. Currents were normalized for cell capacitance and expressed in pA/pF. Membrane potential was recorded under the current-clamp mode.

### Statistical analysis

All data are presented as the mean ± standard error of the mean (SEM). Two-tailed Student’s *t*-test was used to compare two groups and one-way or two-way ANOVA tests followed by Bonferroni’s *post-hoc* test were used when comparing three or more groups, according with the conditions of normality and homoscedasticity. Shapiro–Wilk and Brown–Forsythe tests were performed to analyze these conditions. In case the assumptions of normality and homoscedasticity were not accomplished, non-parametrical Kruskal–Wallis test followed by Dunn’s *post-hoc* test was used to compare three or more groups, or two-tailed Student’s *t*-test with Welch’s correction when comparing two groups. A *P*-value <0.05 was considered significant. Additionally, when linear trend tests were performed, *P-*trend <0.05 was considered significant. All of the statistical analyses were performed on GraphPad Prism 7.0a software (San Diego, CA, USA).

## RESULTS

### CSE exposure diminishes hPAFib and hPASMC proliferative capacity

Vascular remodeling during PH is due to hypertrophy and/or hyperplasia of the predominant cell types within each of the three layers in pulmonary arteries, these including fibroblasts, smooth muscle cells and endothelial cells [40]. Studies in animal models and human cells demonstrate a wide variability of results with respect to the effects of CSE in cellular proliferation. In human airway structural cells a pro-proliferative effect has been observed [12, 41, 42], but opposite results were observed in human pulmonary artery endothelial cells [17]. However, the direct effects of CSE on human pulmonary artery fibroblasts and smooth muscle cells have not been evaluated. To clarify if the effects of CSE on pulmonary cell proliferation depend on different cell lung compartments, we assessed the effects of CSE on proliferation of human pulmonary arterial cells such as hPASMC and hPAFib. To this aim, we stimulated the cells for 24h with increasing concentrations of CSE trapped in aqueous solution and containing the volatile components of cigarettes. Cell proliferation was evaluated using CellTrace™ CFSE and quantifying the loss of CFSE fluorescence by flow cytometry. Our results proved that untreated hPAFib (Figure 1, upper left panel) and hPASMC (Figure 1, upper right panel) showed a significant loss of CFSE fluorescence compared to 10% CSE-treated cells, which indicated a decreased proliferation rate in the treated cells. Quantitative analysis indicated that CSE exposure decreased cell proliferation in a dose-dependent manner compared with untreated cells. This response reached statistical significance from a concentration of 5% CSE with a maximum decrease at 15% CSE (Figure 1, lower panels). Of note, concentrations above the used ones induced cell death (data not shown).

**Figure 1.**
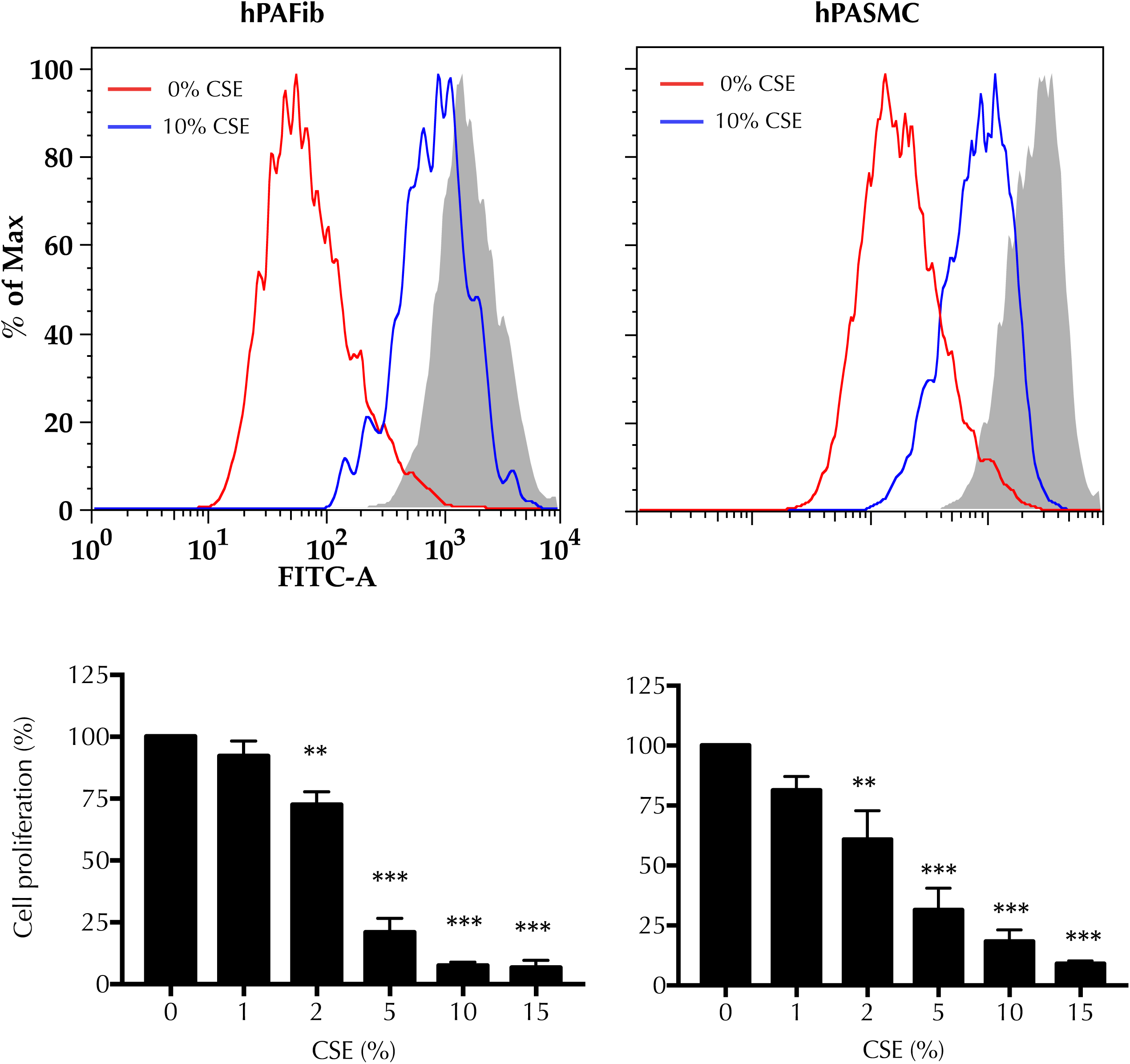
CSE exposure diminishes hPAFib and hPASMC proliferative capacity. CFSE dilution assays were performed to evaluate changes in hPAFib and hPASMC proliferation after a 72-hour challenge to increasing CSE amounts. Representative overlay histograms of CFSE-labelled hPAFib (left) and hPASMC (right) proliferation are shown: control (red line) and 10% CSE-exposed cells (blue line) compared to day 0 staining (filled gray histogram). Proliferation levels are expressed as the decrease in median CFSE fluorescence with respect to day 0 and as control-normalized percentages. Data are presented as mean±SEM. Statistical comparisons between different conditions were made using one-way ANOVA test followed by Bonferroni’s *post-hoc* test (\*\**P*<0.01, \*\*\**P*<0.005), n=4.

### Cellular senescence associated markers are increased after acute CSE treatment in hPAFib and hPASMC

Previous reports in the literature have shown a dual proliferative and senescent phenotype in arteries of COPD patients [18]. In addition, results from *in vitro* studies indicate that CSE induces a senescent phenotype in pulmonary artery endothelial cells, lung parenchyma fibroblasts and airway cells [15, 17, 43]. Taking into account our results above indicating that CSE treatment decreased proliferation of hPAFib and hPASMC, we hypothesized that CSE might induce senescence in these cells. Increased levels of the two main senescence pathways, p53-p21 and p16-retinoblastoma protein (pRb) [44], along with increased activity of β-galactosidase (β-Gal) and cell hypertrophy are hallmarks for cellular senescence [45]. Therefore, we analyzed these markers in hPAFib and hPASMC exposed to different concentrations of CSE. Our results showed that CSE treated cells exhibited a phenotype that is typical of cellular senescence, with a CSE-concentration-dependent increase in β-Gal activity (Figure 2A). This increase was clearly marked in hPAFib and statistically significant at 10% CSE and above and showed a clear upward trend in hPASMC (Figure 2A). In addition, the cells displayed a characteristic enlarged morphology (Figure 2B).

**Figure 2.**
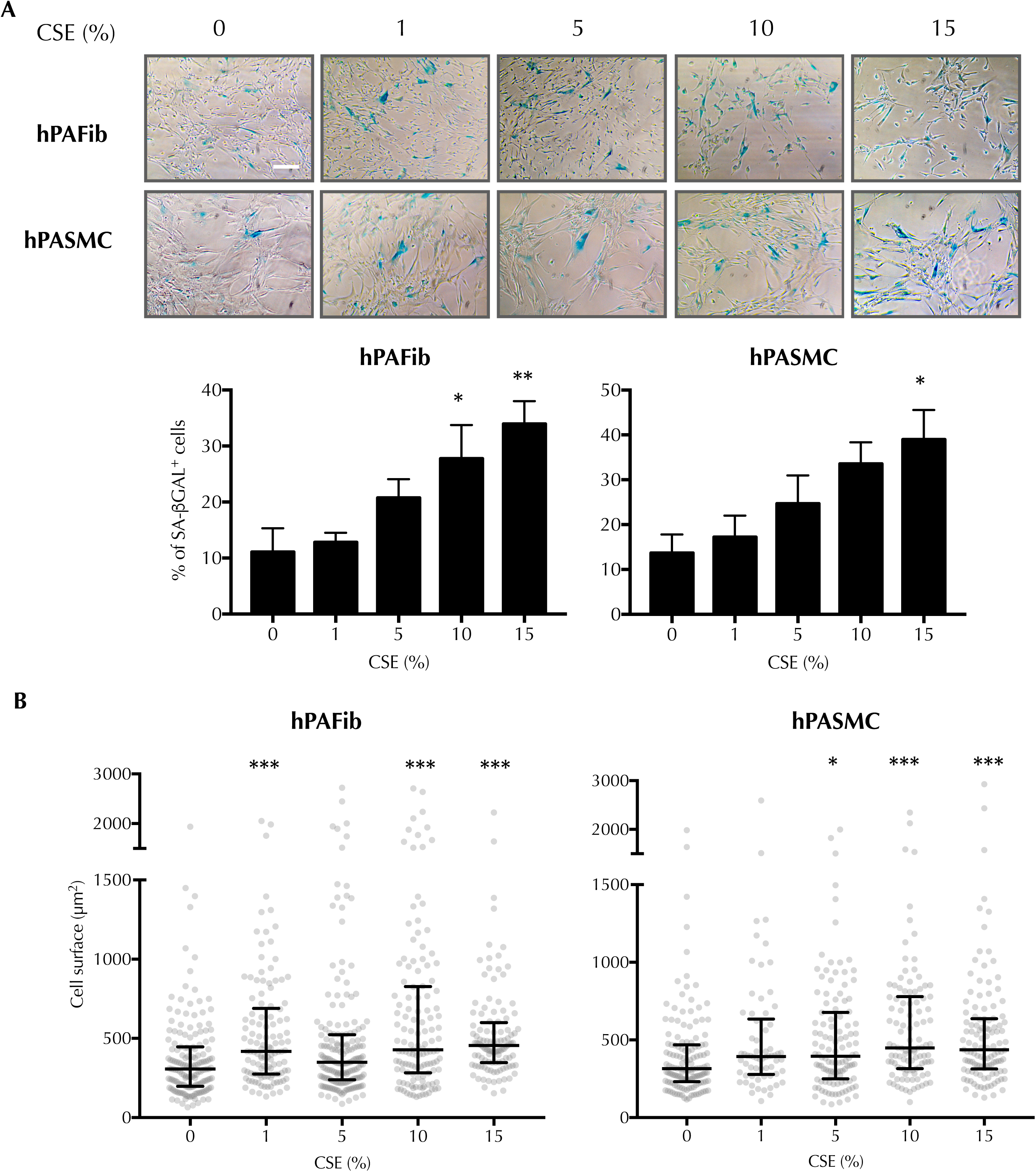
Cellular senescence associated markers are increased after acute CSE treatment in hPAFib and hPASMC. (A) X-Gal staining was used to analyze SA-β-galactosidase activity in hPAFib and hPASMC following CSE-exposure for 48 hours. Representative phase contrast images are shown, scale bar = 100 µm. Data are presented as percentage of blue-stained cells over total cells on three random fields per condition, and expressed as mean±SEM. Statistical comparisons between different conditions were made using one-way ANOVA test followed by Bonferroni’s *post-hoc* test (\**P*<0.05, \*\**P*<0.01), n=5. (B) CSE-induced hypertrophy was assessed by measuring apparent cell surface after 24- hour challenge. Black lines represent median and interquartile range of each population. Statistical comparisons between different conditions were made using Kruskal–Wallis test followed by Dunn’s *post-hoc* test (\**P*<0.05, \*\*\**P*<0.005). More than 100 cells per condition from three independent experiments were analyzed.

Next, we analyzed the expression of the senescence markers p53, p21 and p16 in hPAFib and hPASMC exposed to increasing CSE concentrations. Analysis of p21 (*CDKN1A)* and p16 (*CDKN2A)* gene expression indicated a significant increase of p21 in hPAFib at the highest CSE dose, and a clear upward trend in hPASMC, while no significant changes in p16 were observed (Figure 3A). Furthermore, the increase in gene expression was accompanied by a significant increase in p21 protein levels in both cell types that was CSE concentration-dependent (Figure 3B). Similarly, we observed a CSE concentration-dependent increase in p53 protein levels in both cell types (Figure 3B). Surprisingly, p16 protein levels decreased after CSE treatment in both cell types (Figure 3B).

**Figure 3.**
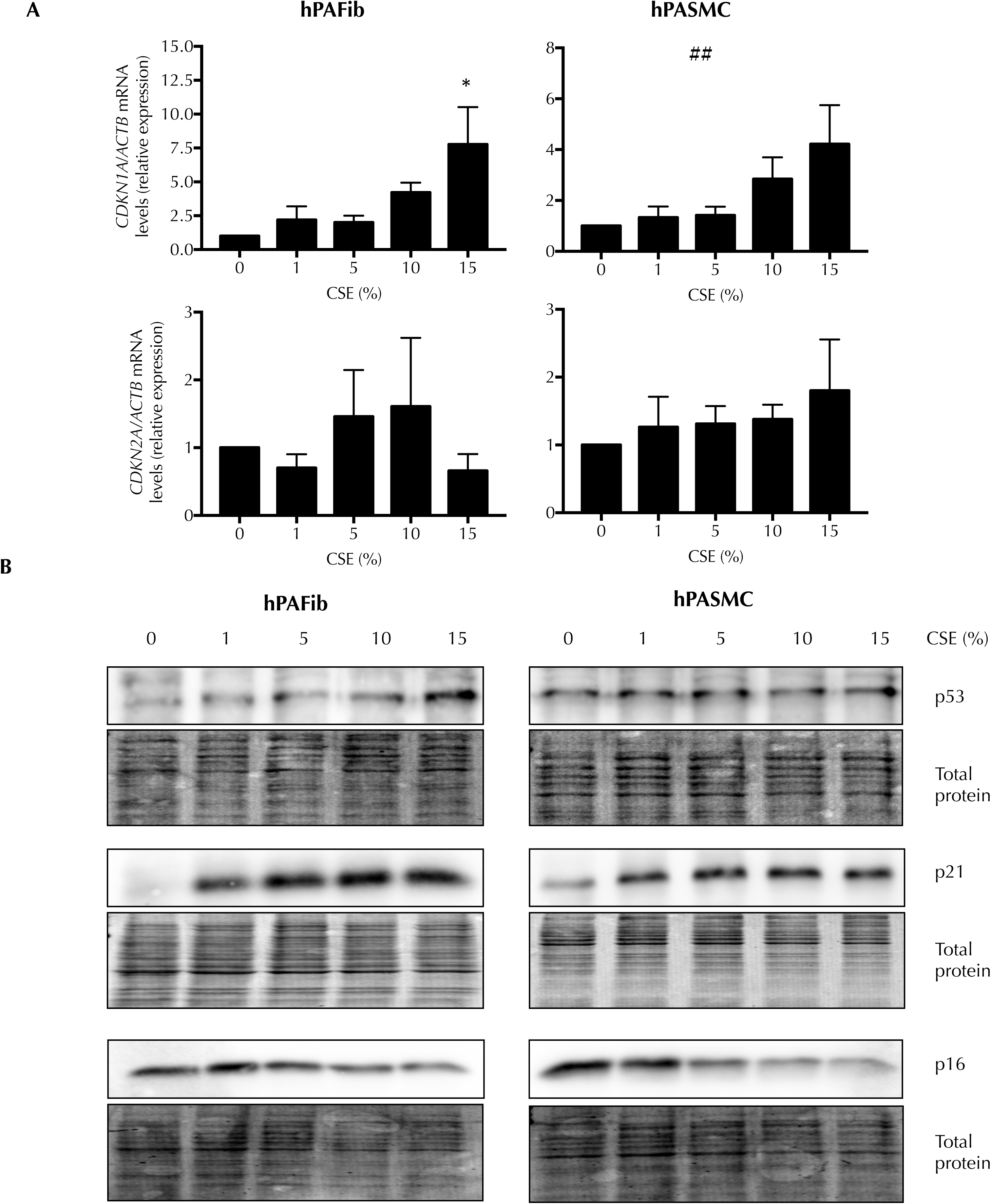
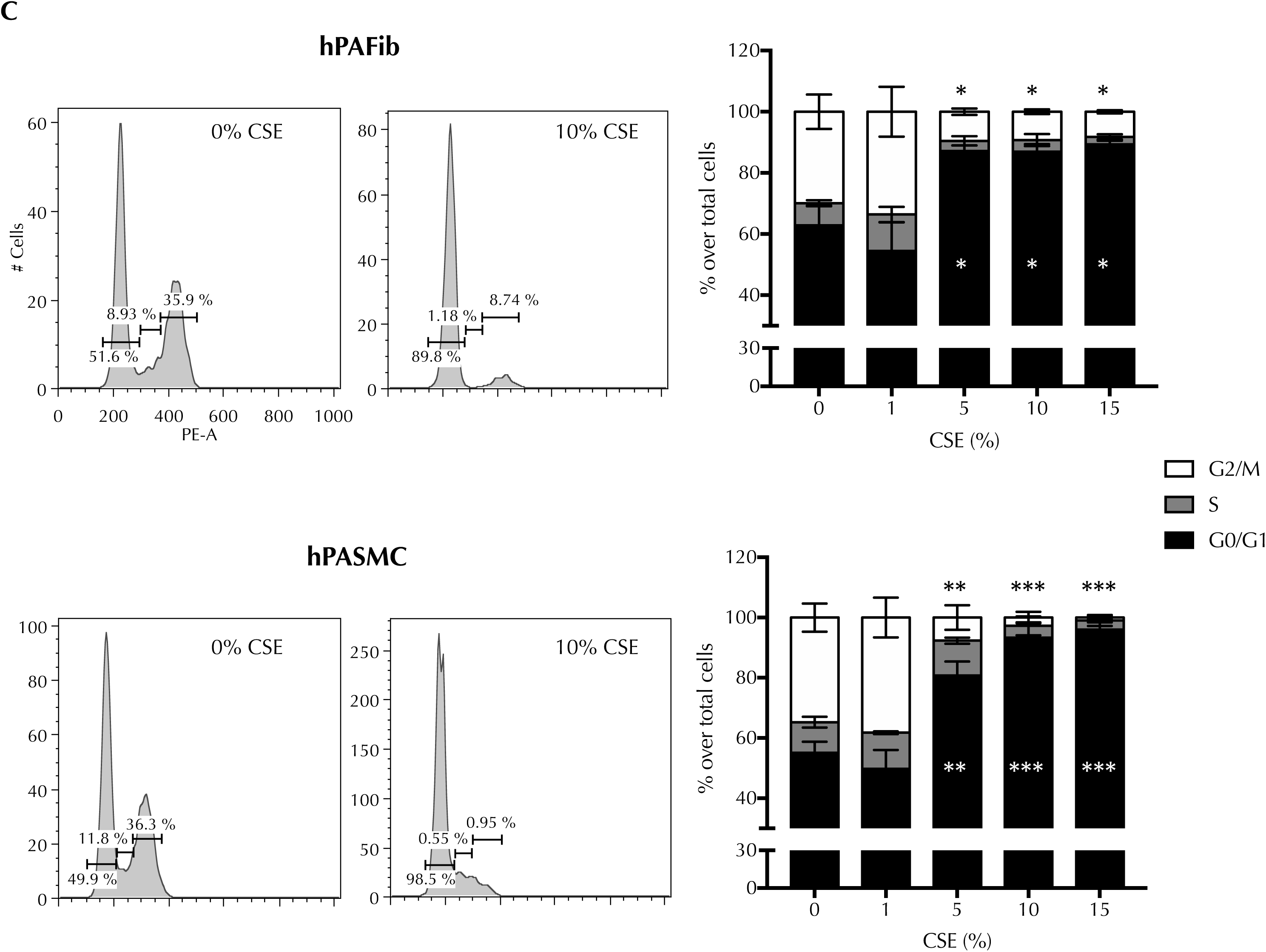
Cellular senescence associated markers are increased after acute CSE treatment in hPAFib and hPASMC. (A) Quantitative RT–PCR analysis was performed to determine *CDKN1A* and *CDKN2A* mRNA expression levels from hPAFib and hPASMC for the indicated CSE concentrations. mRNA levels are expressed as fold change over control cells, controlled with *ACTB* as housekeeping gene and presented as mean±SEM, n=3. (B) Protein levels from hPAFib and hPASMC exposed to increasing amounts of CSE for 24 hours were detected by western blot probed against p53, p21, p16 and total protein as loading control. Representative images of three experiments are shown. (C) Representative propidium iodide DNA-staining histograms showing cell cycle distribution of hPAFib and hPASMC exposed to increasing CSE concentrations for 24 hours. Data are expressed as percentage of cells at each cell cycle phase over total cells exposed to increasing CSE concentration for 24 hours and presented as mean±SEM. Statistical analysis was carried out using one-way ANOVA test followed by Bonferroni’s *post-hoc* test (\**P*<0.05, \*\**P*<0.01, \*\*\**P*<0.005, ^##^*P-trend*<0.01), n=3.

Since cell proliferation is related to cell cycle changes and p21 and p16 are well-known cyclin-dependent kinase inhibitors [46], we further investigated the effect of CSE treatment on cell cycle progression by analyzing propidium iodide incorporation in hPAFib and hPASMC. Our results proved that cells treated with increasing concentrations of CSE underwent a clear arrest in G0/G1 phase compared with untreated cells (Figure 3C).

### CSE-mediated secretion of pro-inflammatory cytokines in hPAFib and hPASMC promotes proliferation of non-exposed cells

Cellular senescence constitutes a strong anti-proliferative response. Interestingly, previous studies demonstrated that senescent PASMC from COPD patients stimulate the growth and migration of healthy PASMC through the release of inflammatory factors such as IL-6, IL-8 and MCP-1[18]. To determine whether CSE-mediated senescence in hPAFib and hPASMC also stimulates a secretory phenotype we analyzed the expression of these factors after acute CSE exposure. Our results showed a significant increase of IL-8 in hPASMC and a similar tendency of IL-6 and IL-8 in hPAFib (Figure 4A), while no changes in the levels of MCP-1 were observed (data not shown). Next, we assessed if conditioned medium from CSE-treated cells containing these soluble factors had paracrine or autocrine effects stimulating proliferation of non-exposed hPAFib and hPASMC. To this aim, we cultured our cells with conditioned media from hPAFib or hPASMC treated with different concentrations of CSE. As shown in figure 4B, conditioned media from both hPAFib and hPASMC stimulated proliferation of non-exposed hPAFib and hPASMC. Interestingly, the effects of conditioned media from hPAFib on cell proliferation were stronger compared to that from hPASMC (Figure 4B).

**Figure 4.**
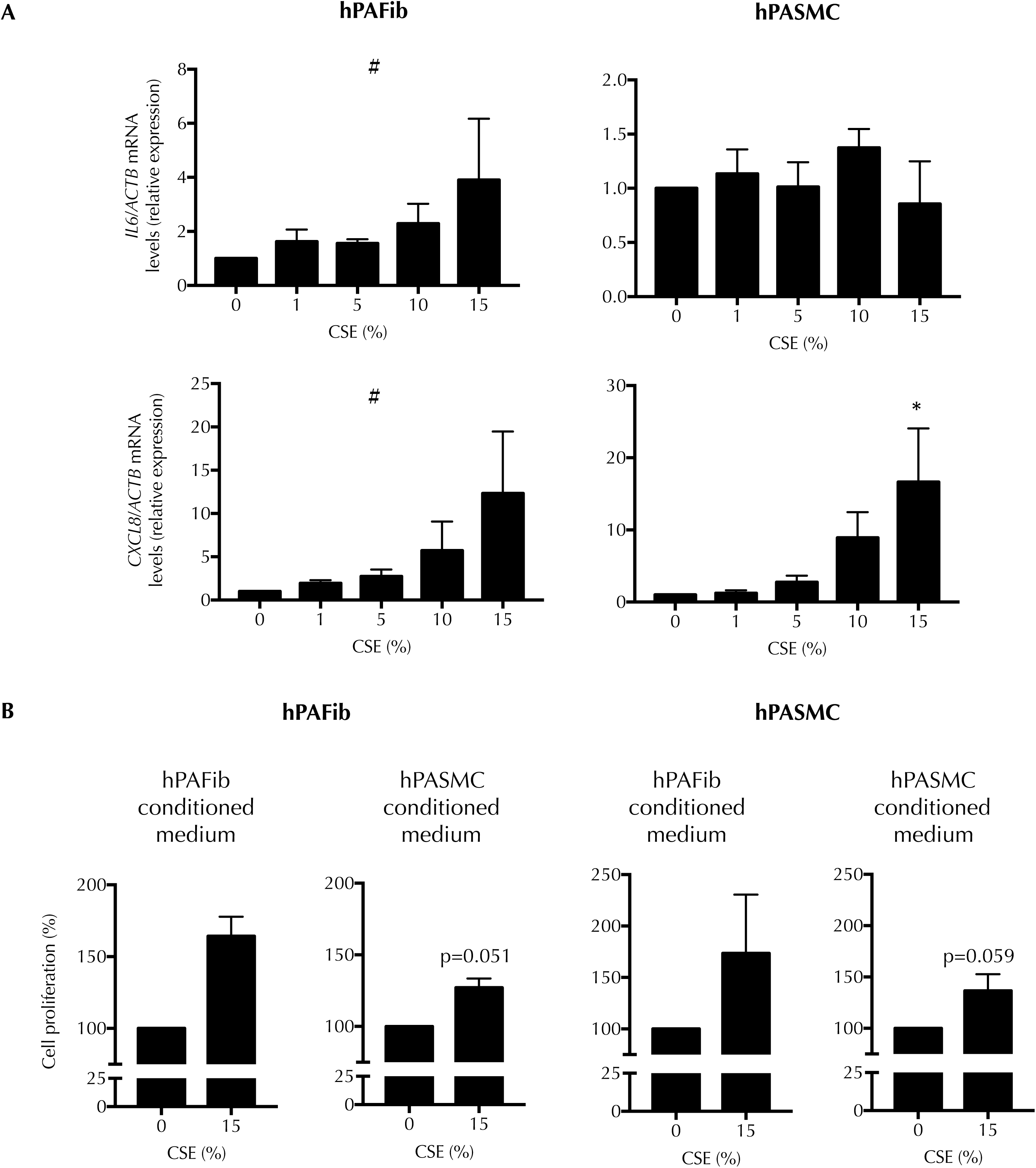
CSE-mediated secretion of pro-inflammatory cytokines in hPAFib and hPASMC promotes proliferation of non-exposed cells. (A) Quantitative RT–PCR analysis was performed to determine *IL6* and *CXCL8* mRNA levels in hPAFib and hPASMC after 24-hour CSE-challenge. mRNA levels are expressed as fold change over control cells, controlled with *ACTB* as housekeeping gene and presented as mean±SEM. Statistical analysis between groups was carried out using one-way ANOVA test followed by Bonferroni’s *post-hoc* test (\**P*<0.05), n=4. (B) CFSE dilution assays were performed to evaluate changes in hPAFib and hPASMC proliferation after a 72-hour challenging to conditioned media from control or CSE-treated hPAFib and hPASMC. Statistical analysis was carried out using two-tailed Student’s t-test with Welch’s correction, n=2.

### CSE increases total ROS levels in hPAFib or hPASMC and upregulates NADPH oxidase subunits NOX1 and p22^**phox**^

We assessed if CSE treatment altered the production of ROS in hPAFib and hPASMC. Analysis by the DHE probe demonstrated that CSE led to an increase in total reactive oxygen species (ROS) production in both hPAFib and hPASMC. This increase was concentration-dependent and reached significance at the highest CSE concentrations used (Figure 5A). Since the nicotinamide adenine dinucleotide phosphate oxidase (NADPH oxidase) complex is considered a major source of ROS production in the pulmonary vasculature [47], we measured gene expression levels of several components of the NADPH oxidase complex. We analyzed the expression of NOX, in particular NOX1 and NOX4, which are expressed in smooth muscle cells [48], and *CYBA* (encoding for the protein p22^phox^). CSE-treated hPAFib and hPASMC showed a concentration-dependent increase of total ROS levels compared with untreated cells (Figure 5A). In addition, the increase in ROS correlated with a significantly higher expression of NOX1 in hPAFib and p22^phox^ in hPASMC (Figure 5B), though we observed the same increasing tendency in both cell types.

**Figure 5.**
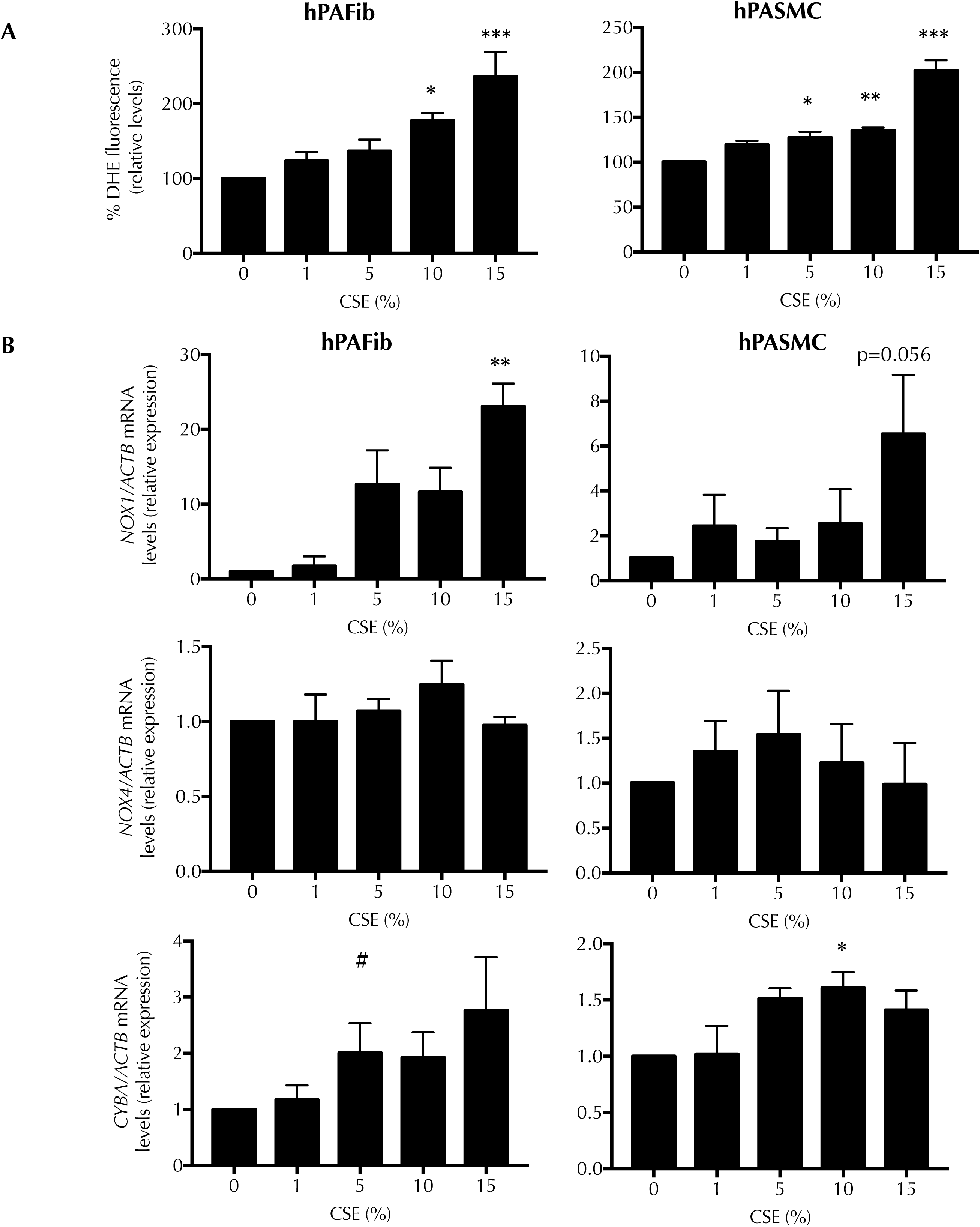
CSE increases total ROS levels in hPAFib and hPASMC and upregulates NADPH oxidase subunits NOX1 and p22phox. (A) Following 24- hour CSE-challenge, 10 µM DHE probe was used to quantify total ROS levels in hPAFib and hPASMC by flow cytometry. Values are calculated as median DHE- treated cells fluorescence minus median vehicle-treated cells fluorescence, presented as control-normalized percentages and expressed as mean±SEM. (B) Quantitative RT–PCR analysis was performed to determine *NOX1, NOX4* and *CYBA* mRNA levels in hPAFib and hPASMC after 24-hour CSE-exposure. mRNA levels are expressed as fold change over control cells, controlled with *ACTB* as housekeeping gene and presented as mean±SEM. Statistical analysis between groups was carried out using one-way ANOVA test followed by Bonferroni’s *post-hoc* test (\**P*<0.05, \*\*\**P*<0.005), n=3.

### Contractile force generation is blocked in hPASMC cells after CSE treatment

PASMC contractile properties are essential for the regulation of blood flow and vascular tone homeostasis throughout the pulmonary system, undergoing constriction under hypoxia in the alveolar compartment [49]. Previous studies highlight the importance of an abnormal mechanical behavior of the pulmonary artery in different pathologies, including COPD and PH [50, 51]. Despite this, the connection between main COPD risk factors, such as cigarette smoke, and biophysical changes in the pulmonary artery cells remains unexplored.

Our results above showed the acquisition of a senescent phenotype induced by CSE-exposure in human PA cells that could be directly responsible for vascular damage and remodeling during COPD. However, it is unclear how the original cell functions are lost in favor of this senescence development, specifically vascular tone regulation and force generation by PASMC. To investigate the direct effects of CSE on the force exerted by hPASMC onto their substrate, we performed 3D-collagen matrix experiments and measured progressive gel contraction along time. After 48-hour CSE challenge, we observed a significant decrease in the surface of gels with control cells that did not occur on gels with 15% CSE-treated cells, indicating a reduction in the total force generation of hPASMC due to cigarette smoke exposure (Figure 6A).

**Figure 6.**
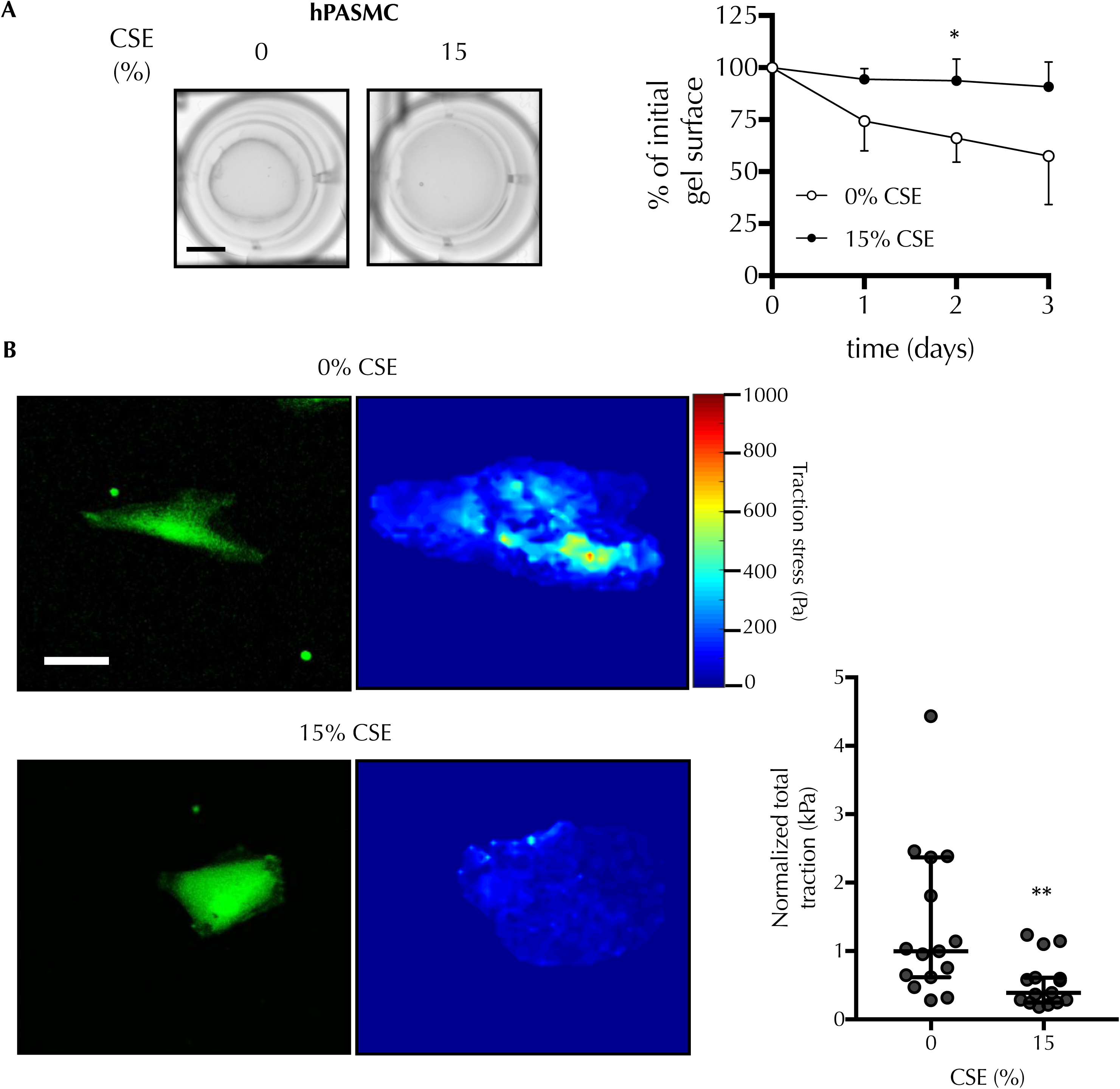
Contractile force generation is blocked in hPASMC after CSE treatment. (A) Contractility assays were performed in 3D collagen matrices to evaluate hPASMC force generation capacity in response to CSE. Representative images of the gels after 3-day treatment are shown, scale bar=0.5 cm. Data are presented as relative percentages of collagen gel surface at the different time points of the treatment, compared to its surface at the beginning of the experiment and expressed as presented as mean±SEM, n=3. (B) TFM experiments were carried out to assess differences in force generation at the single-cell level between control and CSE-exposed hPASMC for 24 hours. Representative examples of confocal images of control and CSE-treated cells (left) and color-coded maps of traction stress magnitude reconstructions (right) are shown, scale bar=15 µm. Quantification of TFM results was measured as total traction stress normalized by the apparent cell surface. Black lines represent median and interquartile range of each population, n=3. Statistical comparisons were made using two-tailed Student’s t-test (\**P*<0.05, \*\**P*<0.01).

Measuring force in 3D collagen gels is often a reliable experimental approach since these gels partly mimic real, tridimensional extracellular matrix architecture. It presents, however, the pitfall of measuring the total force exerted by multiple cells and is strongly dependent on the number of cells embedded within the gel. Our results precisely showed a significant decrease on cell proliferation after CSE treatment that could also explain the diminished force exerted by hPASMC on collagen matrices. To avoid the interference of senescent cells, we turned to traction force microscopy experiments in order to quantify forces on individualized cells. These experiments demonstrated a significant decrease in traction stress force also at the single-cell level exerted on the substrate when hPASMC were exposed to 15% CSE (Figure 6B and Supplementary Figure S2).

### CSE impairs vasoactive responses and induces muscle cell depolarization in murine pulmonary arteries

Since we found a significant decrease in the traction force potential of CSE-treated hPASMC, we next asked whether this effect would also translate into the mechanics of the whole pulmonary artery. To that end, we dissected intact PAs from male wild type mice and exposed them to CSE in order to evaluate their contractility responses on a wire myograph. We first observed a significantly lower contractile response to KCl in CSE-pretreated PAs compared to untreated (Figure 7A), suggesting a general impairment in vascular contractility due to CSE. After that, we analyzed the effects of more specific vasoactive substances, such as serotonin (5-HT) and acetylcholine (ACh), on control and CSE-challenged PAs. Similarly, to the effects of extracellular K^+^, serotonin-induced vasoconstriction (Figure 7B) was significantly decreased in CSE-challenged PAs compared to controls.

**Figure 7.**
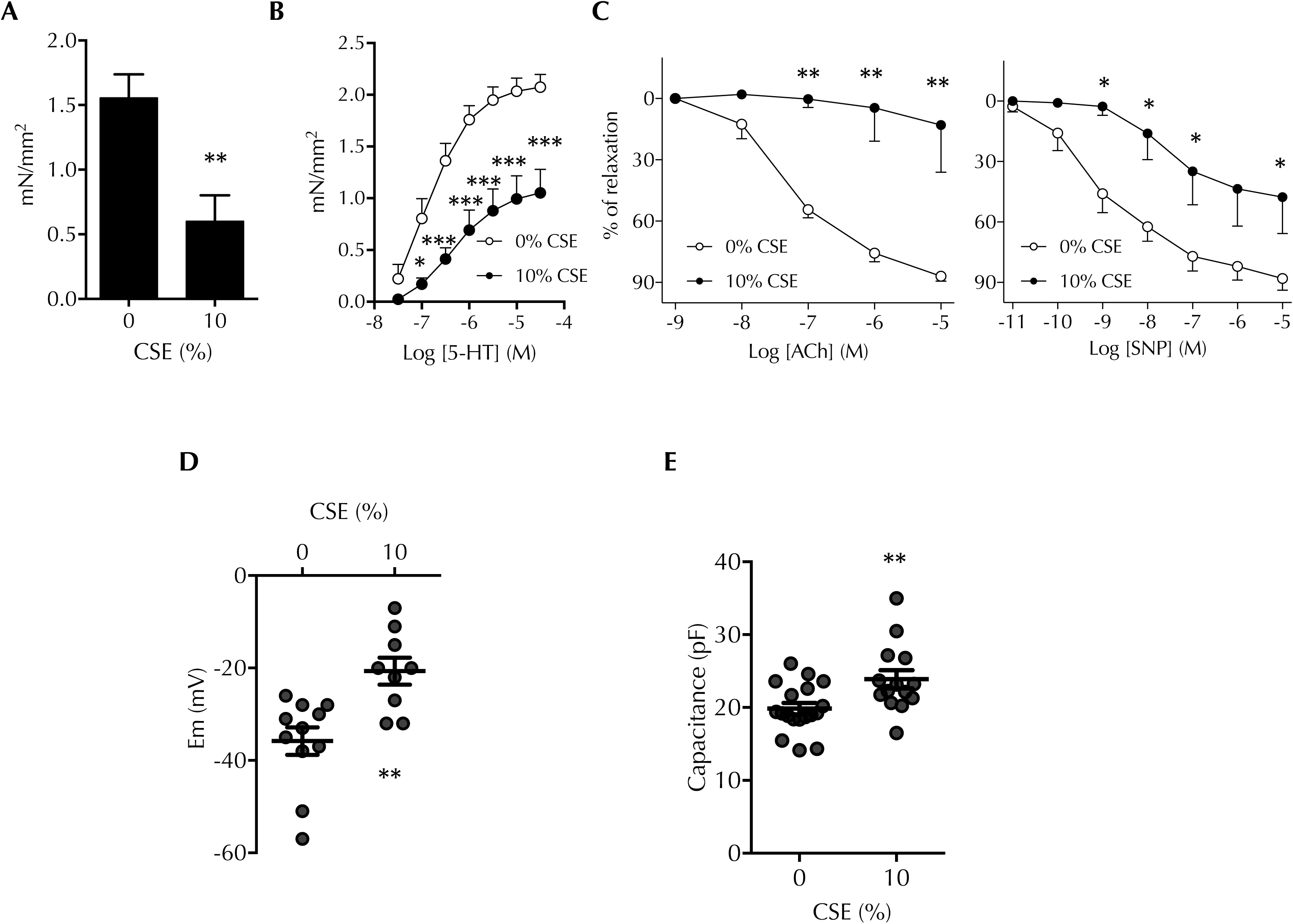
CSE impairs vasoactive responses and induces muscle cell depolarization in murine pulmonary arteries. Vascular responses were analyzed in endothelium-intact PAs from WT mice previously incubated with CSE overnight. Average values of the vasoconstriction in 80 mM KCl (A) or 10 µM serotonin (5-HT) stimulated PAs (B). (C) Average values of the vasodilation induced by increasing amounts of acetylcholine (ACh) (left) or sodium nitroprusside (SNP) (right) following 5-HT stimulation. (D) Resting membrane potential of SMC from control or 10% CSE-challenged PAs, measured under current clamp mode. (E) Average cell capacitance of SMC from control or 10% CSE-challenged PAs. Data are presented as mean±SEM, n=7 (A-C), or as median and interquartile range of each population, n=7 (0% CSE), n=5 (10% CSE) (D-E). Statistical comparisons were made using one-way (A), two-way ANOVA (B-C) followed by Bonferroni’s *post-hoc* test or two-tailed Student’s t-test (D-E) (\**P* < 0.05, \*\**P*<0.01, \*\*\**P*<0.005).

Moreover, the endothelium-dependent relaxation induced by acetylcholine was nearly blunted in CSE-exposed PA, indicating that CSE induces endothelium dysfunction (Figure 7C, left). We also studied the effects of sodium nitroprusside (SNP), an endothelium-independent NO donor. As shown in Figure 7C (right), vasodilation capacity of CSE-challenged PAs in response to SNP was significantly lower compared to control Pas, suggesting an impaired function of vascular smooth muscle cells as well.

Because SMC are excitable cells, some of the effects we observed on CSE-exposed PAs might be related to changes in the electrical properties of these cells. Following SMC isolation from control or CSE-treated murine PAs, we evaluated their membrane potential by means of whole-cell recording configuration of the patch clamp technique, and observed a significant cell depolarization due to cigarette smoke exposure (Figure 7D). In addition, these cells showed higher membrane capacitance, as an indicator of SMC hypertrophy, when exposed to CSE (Figure 7E). This is in agreement with our results in culture hPASMC shown in Figure 2B.

### Kv7.4 channel levels and functionality are diminished in CSE-exposed pulmonary arteries

Changes in expression and activity of voltage-gated potassium (Kv) channels play a key role in vascular tone deregulation and development of lung vasculopathies, such as pulmonary hypertension. Our results showed a notable depolarization of PA cells in response to CSE exposure, which suggested a misbalance of the ion currents in charge of membrane potential homeostasis. Considering this, we hypothesized that K^+^ channel function could be altered in CSE-challenged cells and be responsible for the vascular alterations described above. When we compared SMC isolated from pretreated or control murine PAs, we observed a significant decrease in K^+^ current density due to cigarette smoke exposure (Figure 8A). Among the different channels contributing to potassium flow homeostasis, Kv1 and Kv7 channels are thought to be especially relevant in vascular pathology. Our results indicated a CSE-concentration-dependent decrease in Kv7.4 protein levels in hPASMC (Figure 8B), while no changes were observed in other K^+^channels such as Kv7.5 or BKCa (Supplementary Figure S3). Interestingly, parallel to Kv7.4 downregulation, levels of Kv1.5, another member of the voltage-gated K^+^ channels family, increased in these cells in response to CSE (Supplementary Figure S3).

**Figure 8.**
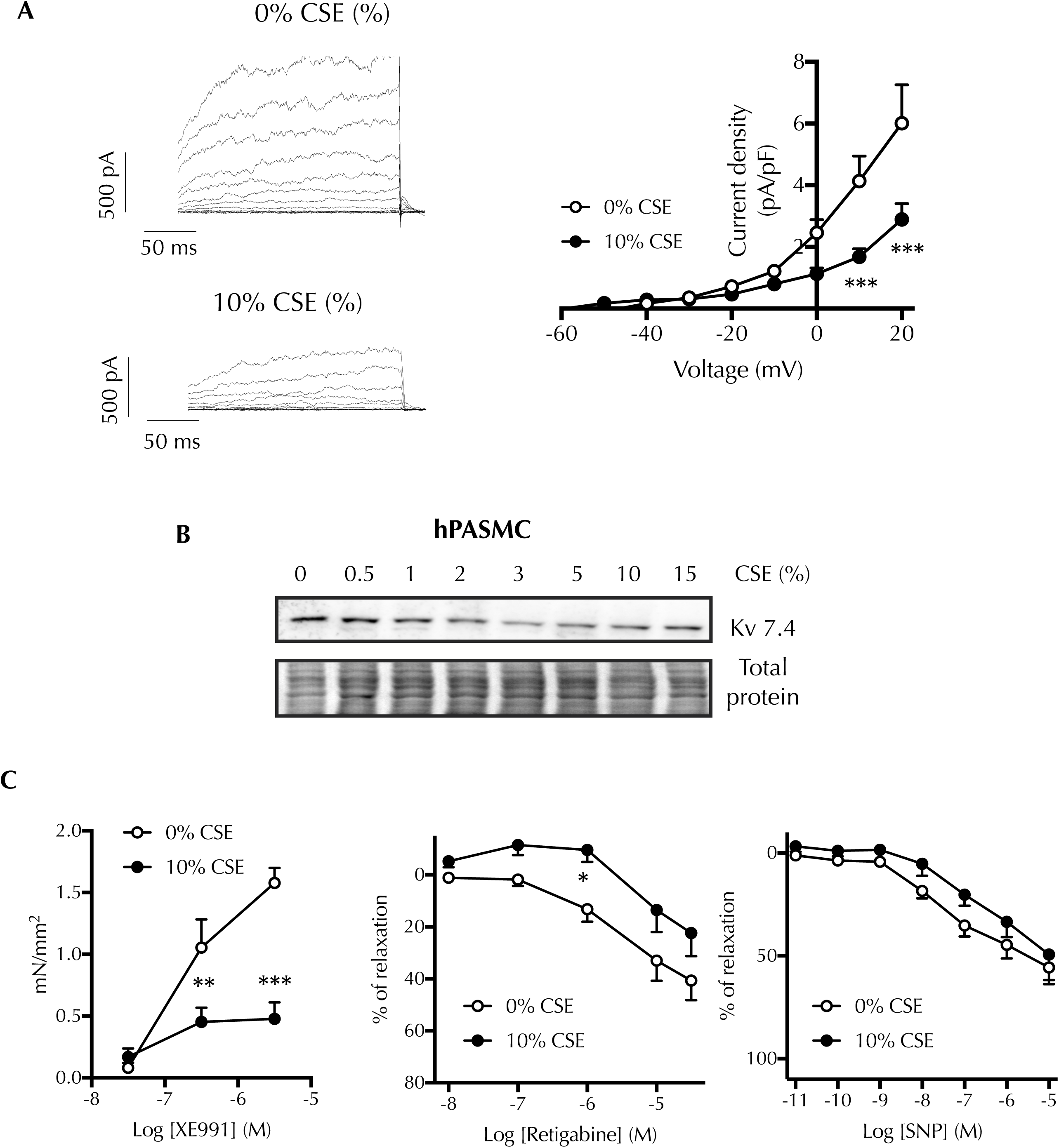
Kv7.4 channel levels and functionality are diminished in CSE-exposed pulmonary arteries. (A) Representative current traces for 200 ms depolarization pulses from −50 mV to +20 mV in 10 mV increments from a holding potential of −50 mV in SMC of control (top) or 10% CSE-exposed PAs. Data are presented as average current-voltage relationships of K^+^ currents, measured at the end of the pulse and normalized by cell capacitance, n=6 (0% CSE), n=5 (10% CSE). (B) Protein levels from hPASMC exposed to increasing amounts of CSE for 24 hours were detected by western blot probed against Kv7.4 and total protein as loading control. (C) Vascular responses were analyzed in endothelium-intact PAs from WT mice previously incubated with 10% CSE overnight. Average values of XE991-induced vasoconstriction (left), retigabine-induced vasodilation (center) and SNP-induced relaxation in presence of XE991 (right) are shown. Data are presented as mean±SEM, n=7. Statistical comparisons were made using two-way ANOVA followed by Bonferroni’s *post-hoc* test (\**P*<0.05, \*\**P*<0.01, \*\*\**P* < 0.005).

To further investigate whether the decrease we observed on Kv7.4 protein levels would also translate into changes on vascular activity, we next analyzed the effect of several Kv7 modulators on control and CSE-exposed murine PAs. The Kv7 inhibitor XE991 induced a contraction which was significantly smaller on PAs exposed to CSE compared to untreated PAs (Figure 8C, left). Furthermore, vasodilation due to the Kv7 activator retigabine was also diminished in CSE-challenged arteries (Figure 8C, center). Importantly, XE991 reduced SNP-dependent relaxation of PAs to practically the same extent in both conditions, so that no statistical differences were found in the presence of this Kv7 channel inhibitor (Figure 8C, right). Together, these results suggested that CSE-treatment also reduced the activity of Kv7 channels, this leading to murine PA vascular dysfunction.

## DISCUSSION

COPD represents an important cause of decreased quality of life and premature death, as well as a risk factor for the presence of other comorbidities. Results from current COPD treatments remain unsatisfactory. Therefore, it is important to understand the cellular and molecular mechanisms that underlie this pathology, to be able to carry out a targeted and effective treatment. In the present study, we identify novel pathogenic factors that could play a role in the development of PH associated to CSE exposure. Our results demonstrate that single CSE exposure had direct effects in the pulmonary vascular bed, and that resident vascular cell types, such as of hPAFib and hPASMC, play specific roles in CSE-mediated remodeling through mechanisms that included cell senescence, cell proliferation, cytokines secretion and ROS production. In addition, our study identifies a K^+^ channel dysfunction by CSE in pulmonary arteries. CSE affected cell contractility and dysregulated the expression and activity of voltage-gated K^+^ channel Kv7.4, this contributing to promote arterial stiffness and limiting vascular responses (proposed model in Figure 9).

**Figure 9.**
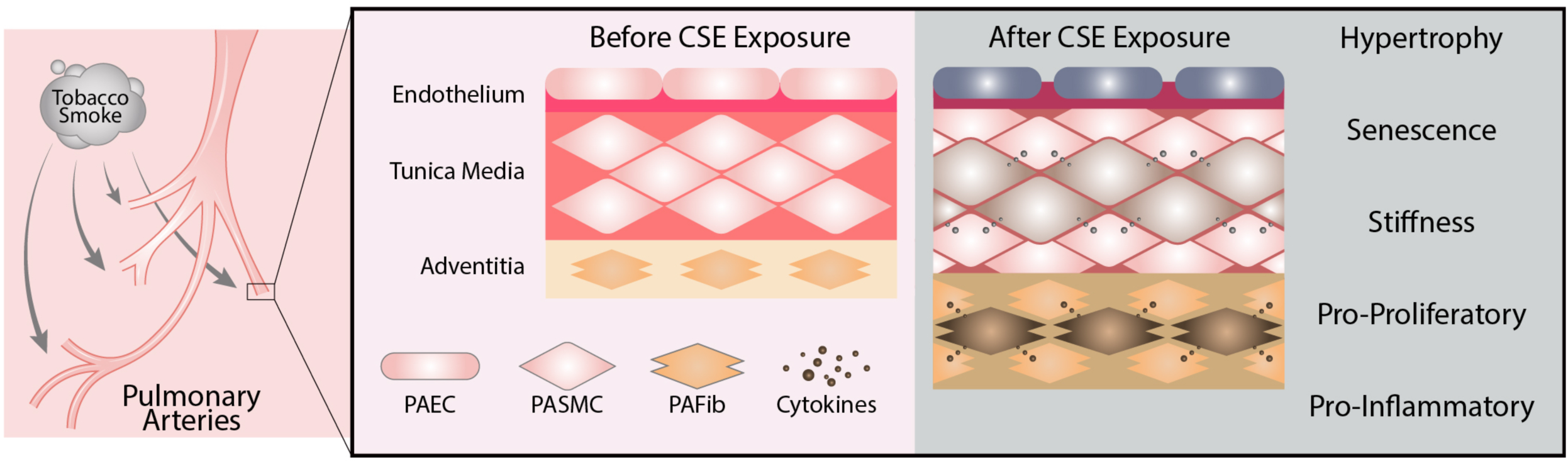
Proposed model: Direct effects of cigarette smoke on pulmonary artery.

The predominant type of pulmonary vascular remodeling in COPD is due to medial hypertrophy, enlargement of the intimal layer in muscular arteries and muscularization of small arterioles [5]. This remodeling process is a contribution of all the cell types in the pulmonary artery that, in response to stress or injury, undergo phenotypic changes and communicate with each other through paracrine or autocrine mechanisms that sustain vascular remodeling. Although cigarette smoke exposure is considered one of the most important agents that contribute to the development of COPD, it is not clear if cigarette smoke effects can be similarly extended to all pulmonary arterial cell types or these are just a consequence of paracrine effects from the airway cells. Studies in both animal models and in human patients have shown that cigarette smoke promotes pulmonary vascular remodeling and pulmonary hypertension development. However, the molecular mechanisms, and whether cigarette smoke have direct effects on the phenotypic changes of pulmonary arterial cells remain unclear. In this paper, we have investigated the consequences of direct effects of CSE on two of the most important cell types during pulmonary arterial remodeling, such as adventitia fibroblasts and media smooth muscle cells. Stenmark and colleagues have demonstrated that pulmonary artery adventitial fibroblasts play critical roles in hypoxia-induced pulmonary vascular remodeling, through phenotypic changes including cell proliferation, migration and recruitment of inflammatory cells [52]. Our results proved that direct CSE exposure of adventitia fibroblasts and media smooth muscle cells contributed to vascular remodeling through various mechanisms, such as induction of a cell senescent phenotype. Conversely, previous reports demonstrate that in human airway structural cells, cigarette smoke exposure promotes a pro-proliferative effect [12, 41, 42]. In agreement with our results, other studies indicate that interstitial lung fibroblasts from COPD patients have a reduced proliferation rate [53]. In addition, results from *in vitro* studies indicate that CSE induces a senescent phenotype in pulmonary artery endothelial cells, lung parenchyma fibroblasts and airway cells [15, 17, 43]. This senescent phenotype in endothelial cells contributes, in a paracrine manner, to stimulate proliferation of PASMC [17]. In the same line of results, other authors have shown an increased number of senescent cells in the lung artery of patients with COPD [18, 54]. Furthermore, PASMC isolated from COPD patients contribute to vascular remodeling through autocrine effects on surrounding PASMC in the media [18]. These results are in agreement with our own results in which we observed that direct CSE effects on PASMC and in PAFib stimulated a secretory phenotype that induced paracrine and autocrine proliferation in non-exposed cells. Interestingly, conditioned media from PAFib showed stronger effects on cell proliferation, this suggesting that these cells might be a more relevant sensor and executor of CSE effects on the artery. In addition, these effects could be mediated through the increased secretion of IL-6 and IL-8 cytokines in the cells. In this respect, Noureddine et al. observed that neutralizing antibodies against IL-6 or MCP-1, but not against IL-8, diminished proliferation of PASMC induced by senescent PASMC conditioned media from COPD patients [18].

Another possibility is that these effects were mediated by the increased ROS levels after CSE treatment. Accumulating data have shown that oxidative stress plays important roles in the pathological changes in the pulmonary vascular bed [55, 56]. In addition, oxidative stress participates in the emergence of cell senescence, in COPD [57]. NADPH oxidases (NOX) are expressed in the vascular wall and are one of the major sources of ROS in the vasculature regulating diverse functions such as proliferation, migration, differentiation, apoptosis, senescence, contractility and inflammatory responses [58-60]. Our results proved a significant increase in the levels of ROS, Nox-1 and p22^phox^ in PASMC and PAFib after CSE treatment. Both, Nox-1 and Nox-4 isoforms are particularly important in the vasculature. Few studies have addressed the role of Nox-4 in COPD, demonstrating that Nox-4 expression and ROS generation are enhanced following exposure of human bronchial epithelial cells to CSE, and this increase correlates with COPD severity [61, 62]. However, we found no significant differences in Nox-4 levels in our pulmonary arterial cells. This indicates that Nox-4 functions in the lung might depend on its cellular compartmentalization in this organ. On the other hand, Nox-1 expression is induced in neointimal SMC after vascular injury [63, 64] where it mediates cell migration, proliferation, and extracellular matrix production, thus contributing to neointima hyperplasia [65], and it is increased in COPD patients [66]. In addition, other authors demonstrated that Nox-1 mediates CSE-induced ROS in rat aortic smooth muscle cells [48]. The increase of the regulatory subunit p22^phox^ observed in our cells might also contribute to the increased activity and stability of this NADPH complex. In this respect, Nagaraj *et al.* demonstrated the crucial role of p22^phox^ in COPD associated pulmonary hypertension, since the lack of p22^phox^ is associated with decreased pulmonary vascular remodeling and improved right ventricular function [67]. Thus, our results contribute to support these previous findings in that they prove a CSE-mediated increase of Nox-1 and p22^phox^ in pulmonary arterial cells. This increase might contribute to rise ROS levels, induce the senescent phenotype and thus to the vascular remodeling during COPD.

Abnormal contraction of COPD pulmonary arteries accounts for much of the condition, after which vascular remodeling becomes predominant. Previous reports demonstrate that CSE impairs endothelium-dependent relaxation of rabbit aortas [68]. Dysregulation of Kv channels is observed in PASMCs from patients with PH [35] and in PASMC from rats with chronic hypoxia-induced PH [69, 70]. *KCNQ* (*KCNQ1–5*) genes encode a subfamily of voltage-gated K^+^ channels, denoted K_v_7.1-K_v_7.5. These channels play a key role in the control of vascular tone in several blood vessels, including the pulmonary circulation [34, 71-73]. Previous results suggest that Kv7 channels are key regulators of vascular tone and reduced expression and activity of Kv7 channels has been reported in several cardiovascular diseases including essential hypertension [71, 72]. Interestingly, reduced *KCNQ4* expression, encoding for Kv7.4, has been reported in early phases of hypoxia-induced pulmonary hypertension [74]. Furthermore, treatment with the Kv7 channel activator flupirtine prevented the development of hypoxia-induced PH in murine models [75]. Very recently we have shown that activation of Kv7 channels contribute to the pulmonary vasodilation induced by nitric oxide (Mondejar–Parreño et al 2019; in press). In line with this, herein we found that the relaxation to the NO donor SNP was reduced in the presence of the Kv7 channel inhibitor XE991. In spite of the growing evidence of the important role of Kv7 channels in the pulmonary circulation the regulation of these channels during pulmonary hypertension in COPD has not been studied. Our results demonstrate for the first time that CSE exposure decreases the levels and activity of Kv7.4 in PASMC. Furthermore, CSE treatment significantly decreases Kv7-dependent vasodilator response in murine PAs, contributing to vascular dysfunction. Interestingly in the presence of the Kv7 channel inhibitor the relaxation to SNP was similar in CSE-versus unexposed-PA, suggesting that downregulation of Kv7 channel may account for the reduced relaxant response following CSE exposure. Other reports demonstrate that the imbalance in the ratio of other channels like BKCa and Kv1.5 might explain in part the etiology of remodeling and PH in COPD patients [76]. Interestingly, our results prove that BKCa levels in PASMC are not affected by CSE treatment. However, we found a significant increase on Kv1.5 levels in PASMC. This might be a compensatory response to the decrease on Kv7.4 levels after CSE treatment. However, another possibility is that the increase on Kv1.5 levels are not functionally relevant since it can be inhibited by the increase of ROS [77]. Therefore, CSE-mediated vascular dysfunction might be at least partly attributed to the decreased levels and activity of Kv7.4.

In conclusion, our results prove that CSE has differential effects in pulmonary arteries compared to effects described in the airways. Furthermore, we corroborate the idea that CSE-induced vascular remodeling is not just a consequence of alveolar hypoxia and loss of capillary bed but can also be due to direct effects on the vascular bed. Therefore, these findings greatly enhance knowledge about pulmonary diseases, in particular COPD, and open a new line of study related to the exploitation of these mechanisms as therapeutic targets.

## AUTHOR’S CONTRIBUTION

MJC and JSM conceived and designed research; JSM, CFP, G.M.P. and B.B. performed experiments; JSM, CFP, G.M.P, B.B, A.C and MJC analyzed data; JSM, G.M.P, B.B, A.C and MJC interpreted results of experiments; JSM, DLA, G.M.P, B.B and A.C prepared figures; DLA designed Graphical abstract; MJC and JSM drafted the manuscript; MJC, JSM, DLA and A.C. edited and revised the manuscript; All authors approved the final version of manuscript.

### ACKNOWLEGMENTS

This work was supported by grants from the Spanish Government (cofunded by European Regional Development Fund, ERDF/FEDER); PI16/02166, 2017/EEUU/03, and Red Temática de Excelencia en Investigación en Hipoxia (SAF 2017-90794-REDT) to MJC and AC; SAF2016-77222-R and Comunidad de Madrid (B2017/BMD-3727 to AC.

**Supplementary table S1. List of the primer pairs used for gene expression analysis by real-time PCR.**

**Supplementary figure S2. Contractile force generation is blocked in hPASMC cells after CSE treatment.** TFM experiments were carried out to assess differences in force generation at the single-cell level between control and CSE-exposed hPASMC for 24 hours. Representative examples of confocal images of control and CSE-treated cells (left) and vectorial maps of traction stress reconstructions (right) are shown, scale bar = 15 μm.

**Supplementary figure S3. Kv7.4 channel levels and activity are diminished in CSE-exposed pulmonary arteries.** Protein levels from hPASMC exposed to increasing amounts of CSE for 24 hours were detected by western blot probed against Kv7.5, BKCa, Kv1.5 and total protein as loading control.

## REFERENCES

1. Lozano, R., et al., Global and regional mortality from 235 causes of death for 20 age groups in 1990 and 2010: a systematic analysis for the Global Burden of Disease Study 2010. Lancet, 2013. 380(9859): p. 2095–128.

2. Miravitlles, M., et al., Spanish Guidelines for Management of Chronic Obstructive Pulmonary Disease (GesEPOC) 2017. Pharmacological Treatment of Stable Phase. Arch Bronconeumol, 2017. 53(6): p. 324–335.

3. Galie, N., et al., [2015 ESC/ERS Guidelines for the diagnosis and treatment of pulmonary hypertension]. Kardiol Pol, 2015. 73(12): p. 1127–206.

4. Seeger, W., et al., Pulmonary hypertension in chronic lung diseases. J Am Coll Cardiol, 2013. 62(25 Suppl): p. D109–16.

5. Peinado, V.I., et al., Endothelial dysfunction in pulmonary arteries of patients with mild COPD. Am J Physiol, 1998. 274(6 Pt 1): p. L908–13.

6. Bogaard, H.J., Hypoxic pulmonary vasoconstriction in COPD-associated pulmonary hypertension: been there, done that? Eur Respir J, 2017. 50(1).

7. Carlsen, J., et al., Pulmonary arterial lesions in explanted lungs after transplantation correlate with severity of pulmonary hypertension in chronic obstructive pulmonary disease. J Heart Lung Transplant, 2013. 32(3): p. 347–54.

8. Barbera, J.A., Mechanisms of development of chronic obstructive pulmonary disease-associated pulmonary hypertension. Pulm Circ, 2013. 3(1): p. 160–4.

9. Peinado, V.I., S. Pizarro, and J.A. Barbera, Pulmonary vascular involvement in COPD. Chest, 2008. 134(4): p. 808–14.

10. Santos, S., et al., Characterization of pulmonary vascular remodelling in smokers and patients with mild COPD. Eur Respir J, 2002. 19(4): p. 632–8.

11. Seimetz, M., et al., Inducible NOS inhibition reverses tobacco-smoke-induced emphysema and pulmonary hypertension in mice. Cell, 2011. 147(2): p. 293– 305.

12. Guan, P., et al., Cigarette smoke extract promotes proliferation of airway smooth muscle cells through suppressing C/EBP-alpha expression. Exp Ther Med, 2017. 13(4): p. 1408–1414.

13. Guo, T., et al., Downregulation of P16 promotes cigarette smoke extractinduced vascular smooth muscle cell proliferation via preventing G1/S phase transition. Exp Ther Med, 2017. 14(1): p. 214–220.

14. Xing, A.P., et al., Cigarette smoke extract stimulates rat pulmonary artery smooth muscle cell proliferation via PKC-PDGFB signaling. J Biomed Biotechnol, 2012. 2012: p. 534384.

15. Nyunoya, T., et al., Cigarette smoke induces cellular senescence. Am J Respir Cell Mol Biol, 2006. 35(6): p. 681–8.

16. Michaud, S.E., et al., Cigarette smoke exposure impairs VEGF-induced endothelial cell migration: role of NO and reactive oxygen species. J Mol Cell Cardiol, 2006. 41(2): p. 275–84.

17. Cai, L., et al., [The stimulation of human pulmonary artery endothelial cells by cigarette smoke extract contributed to cell senescence and induced human pulmonary artery smooth cell migration]. Zhonghua Jie He He Hu Xi Za Zhi, 2017. 40(6): p. 463–468.

18. Noureddine, H., et al., Pulmonary artery smooth muscle cell senescence is a pathogenic mechanism for pulmonary hypertension in chronic lung disease. Circ Res, 2011. 109(5): p. 543–53.

19. Rode, L., et al., Short telomere length, lung function and chronic obstructive pulmonary disease in 46,396 individuals. Thorax, 2013. 68(5): p. 429–35.

20. Savale, L., et al., Shortened telomeres in circulating leukocytes of patients with chronic obstructive pulmonary disease. Am J Respir Crit Care Med, 2009. 179(7): p. 566–71.

21. Sundar, I.K., et al., Genetic ablation of histone deacetylase 2 leads to lung cellular senescence and lymphoid follicle formation in COPD/emphysema. FASEB J, 2018. 32(9): p. 4955–4971.

22. Even, B., et al., Heme oxygenase-1 induction attenuates senescence in chronic obstructive pulmonary disease lung fibroblasts by protecting against mitochondria dysfunction. Aging Cell, 2018. 17(6): p. e12837.

23. Archer, S.L., et al., Nitric oxide and cGMP cause vasorelaxation by activation of a charybdotoxin-sensitive K channel by cGMP-dependent protein kinase. Proc Natl Acad Sci U S A, 1994. 91(16): p. 7583–7.

24. Yuan, X.J., et al., NO hyperpolarizes pulmonary artery smooth muscle cells and decreases the intracellular Ca2+ concentration by activating voltage-gated K+ channels. Proc Natl Acad Sci U S A, 1996. 93(19): p. 10489–94.

25. Barnes, E.A., et al., Loss of smooth muscle cell hypoxia inducible factor-1alpha underlies increased vascular contractility in pulmonary hypertension. FASEB J, 2017. 31(2): p. 650–662.

26. Burg, E.D., C.V. Remillard, and J.X. Yuan, Potassium channels in the regulation of pulmonary artery smooth muscle cell proliferation and apoptosis: pharmacotherapeutic implications. Br J Pharmacol, 2008. 153 Suppl 1: p. S99-S111.

27. Cogolludo, A., L. Moreno, and E. Villamor, Mechanisms controlling vascular tone in pulmonary arterial hypertension: implications for vasodilator therapy. Pharmacology, 2007. 79(2): p. 65–75.

28. Wang, J., et al., Chronic hypoxia inhibits Kv channel gene expression in rat distal pulmonary artery. Am J Physiol Lung Cell Mol Physiol, 2005. 288(6): p. L1049–58.

29. Yuan, X.J., et al., Attenuated K+ channel gene transcription in primary pulmonary hypertension. Lancet, 1998. 351(9104): p. 726–7.

30. Kazama, I. and T. Tamada, Lymphocyte Kv1.3-channels in the pathogenesis of chronic obstructive pulmonary disease: novel therapeutic implications of targeting the channels by commonly used drugs. Allergy Asthma Clin Immunol, 2016. 12: p. 60.

31. Yu, Z.H., et al., Up-regulation of KCa3.1 promotes human airway smooth muscle cell phenotypic modulation. Pharmacol Res, 2013. 77: p. 30-8.

32. Yu, Z.H., et al., Targeted inhibition of KCa3.1 channel attenuates airway inflammation and remodeling in allergic asthma. Am J Respir Cell Mol Biol, 2013. 48(6): p. 685–93.

33. Boucherat, O., et al., Potassium channels in pulmonary arterial hypertension. Eur Respir J, 2015. 46(4): p. 1167–77.

34. Mondejar-Parreno, G., et al., HIV transgene expression impairs K(+) channel function in the pulmonary vasculature. Am J Physiol Lung Cell Mol Physiol, 2018. 315(5): p. L711–L723.

35. Yuan, J.X., et al., Dysfunctional voltage-gated K+ channels in pulmonary artery smooth muscle cells of patients with primary pulmonary hypertension. Circulation, 1998. 98(14): p. 1400–6.

36. Plotnikov, S.V., et al., High-resolution traction force microscopy. Methods Cell Biol, 2014. 123: p. 367–94.

37. Martiel, J.L., et al., Measurement of cell traction forces with ImageJ. Methods Cell Biol, 2015. 125: p. 269–87.

38. Tseng, Q., et al., Spatial organization of the extracellular matrix regulates cell-cell junction positioning. Proc Natl Acad Sci U S A, 2012. 109(5): p. 1506–11.

39. Cogolludo, A., et al., Serotonin inhibits voltage-gated K+ currents in pulmonary artery smooth muscle cells: role of 5-HT2A receptors, caveolin-1, and KV1.5 channel internalization. Circ Res, 2006. 98(7): p. 931–8.

40. Jeffery, T.K. and J.C. Wanstall, Pulmonary vascular remodeling: a target for therapeutic intervention in pulmonary hypertension. Pharmacol Ther, 2001. 92(1): p. 1–20.

41. He, F., et al., The pro-proliferative effects of nicotine and its underlying mechanism on rat airway smooth muscle cells. PLoS One, 2014. 9(4): p. e93508.

42. Pera, T., et al., Cigarette smoke and lipopolysaccharide induce a proliferative airway smooth muscle phenotype. Respir Res, 2010. 11: p. 48.

43. Tsuji, T., K. Aoshiba, and A. Nagai, Cigarette smoke induces senescence in alveolar epithelial cells. Am J Respir Cell Mol Biol, 2004. 31(6): p. 643–9.

44. Campisi, J., Senescent cells, tumor suppression, and organismal aging: good citizens, bad neighbors. Cell, 2005. 120(4): p. 513–22.

45. Rodier, F. and J. Campisi, Four faces of cellular senescence. J Cell Biol, 2011. 192(4): p. 547–56.

46. Biggs, J.R. and A.S. Kraft, Inhibitors of cyclin-dependent kinase and cancer. J Mol Med (Berl), 1995. 73(10): p. 509–14.

47. Weir, E.K., et al., Acute oxygen-sensing mechanisms. N Engl J Med, 2005. 353(19): p. 2042–55.

48. Chang, K.H., et al., NADPH oxidase (NOX) 1 mediates cigarette smoke-induced superoxide generation in rat vascular smooth muscle cells. Toxicol In Vitro, 2016. 38: p. 49–58.

49. Harkness, L.M., et al., Pulmonary vascular changes in asthma and COPD. Pulm Pharmacol Ther, 2014. 29(2): p. 144–55.

50. Agoston-Coldea, L., S. Lupu, and T. Mocan, Pulmonary Artery Stiffness by Cardiac Magnetic Resonance Imaging Predicts Major Adverse Cardiovascular Events in patients with Chronic Obstructive Pulmonary Disease. Sci Rep, 2018. 8(1): p. 14447.

51. Zambrano, B.A., et al., Image-based computational assessment of vascular wall mechanics and hemodynamics in pulmonary arterial hypertension patients. J Biomech, 2018. 68: p. 84–92.

52. Stenmark, K.R., et al., Role of the adventitia in pulmonary vascular remodeling. Physiology (Bethesda), 2006. 21: p. 134–45.

53. Holz, O., et al., Lung fibroblasts from patients with emphysema show a reduced proliferation rate in culture. Eur Respir J, 2004. 24(4): p. 575–9.

54. Tsuji, T., K. Aoshiba, and A. Nagai, Alveolar cell senescence in patients with pulmonary emphysema. Am J Respir Crit Care Med, 2006. 174(8): p. 886–93.

55. Voelkel, N.F., et al., Antioxidants for the treatment of patients with severe angioproliferative pulmonary hypertension? Antioxid Redox Signal, 2013. 18(14): p. 1810–7.

56. Wong, C.M., et al., Reactive oxygen species and antioxidants in pulmonary hypertension. Antioxid Redox Signal, 2013. 18(14): p. 1789–96.

57. Barnes, P.J., Senescence in COPD and Its Comorbidities. Annu Rev Physiol, 2017. 79: p. 517–539.

58. Lassegue, B., A. San Martin, and K.K. Griendling, Biochemistry, physiology, and pathophysiology of NADPH oxidases in the cardiovascular system. Circ Res, 2012. 110(10): p. 1364–90.

59. Garcia-Redondo, A.B., et al., NADPH oxidases and vascular remodeling in cardiovascular diseases. Pharmacol Res, 2016. 114: p. 110–120.

60. Perez-Vizcaino, F., A. Cogolludo, and L. Moreno, Reactive oxygen species signaling in pulmonary vascular smooth muscle. Respir Physiol Neurobiol, 2010. 174(3): p. 212–20.

61. Liu, X., et al., The Expression of NOX4 in Smooth Muscles of Small Airway Correlates with the Disease Severity of COPD. Biomed Res Int, 2016. 2016: p. 2891810.

62. Milara, J., et al., Roflumilast N-oxide inhibits bronchial epithelial to mesenchymal transition induced by cigarette smoke in smokers with COPD. Pulm Pharmacol Ther, 2014. 28(2): p. 138–48.

63. Szocs, K., et al., Upregulation of Nox-based NAD(P)H oxidases in restenosis after carotid injury. Arterioscler Thromb Vasc Biol, 2002. 22(1): p. 21–7.

64. Xu, S., et al., Increased expression of Nox1 in neointimal smooth muscle cells promotes activation of matrix metalloproteinase-9. J Vasc Res, 2012. 49(3): p. 242–8.

65. Lee, M.Y., et al., Mechanisms of vascular smooth muscle NADPH oxidase 1 (Nox1) contribution to injury-induced neointimal formation. Arterioscler Thromb Vasc Biol, 2009. 29(4): p. 480–7.

66. Schneider, D., et al., Increased cytokine response of rhinovirus-infected airway epithelial cells in chronic obstructive pulmonary disease. Am J Respir Crit Care Med, 2010. 182(3): p. 332–40.

67. Nagaraj, C., et al., Hypoxic vascular response and ventilation/perfusion matching in end-stage COPD may depend on p22phox. Eur Respir J, 2017. 50(1).

68. Ota, Y., et al., Impairment of endothelium-dependent relaxation of rabbit aortas by cigarette smoke extract--role of free radicals and attenuation by captopril. Atherosclerosis, 1997. 131(2): p. 195–202.

69. Platoshyn, O., et al., Chronic hypoxia decreases K(V) channel expression and function in pulmonary artery myocytes. Am J Physiol Lung Cell Mol Physiol, 2001. 280(4): p. L801–12.

70. Smirnov, S.V., et al., Chronic hypoxia is associated with reduced delayed rectifier K+ current in rat pulmonary artery muscle cells. Am J Physiol, 1994. 266(1 Pt 2): p. H365-70.

71. Chadha, P.S., et al., Reduced KCNQ4-encoded voltage-dependent potassium channel activity underlies impaired beta-adrenoceptor-mediated relaxation of renal arteries in hypertension. Hypertension, 2012. 59(4): p. 877–84.

72. Jepps, T.A., et al., Downregulation of Kv7.4 channel activity in primary and secondary hypertension. Circulation, 2011. 124(5): p. 602–11.

73. Morales-Cano, D., et al., Kv7 channels critically determine coronary artery reactivity: left-right differences and down-regulation by hyperglycaemia. Cardiovasc Res, 2015. 106(1): p. 98–108.

74. Sedivy, V., et al., Role of Kv7 channels in responses of the pulmonary circulation to hypoxia. Am J Physiol Lung Cell Mol Physiol, 2015. 308(1): p. L48–57.

75. Morecroft, I., et al., Treatment with the Kv7 potassium channel activator flupirtine is beneficial in two independent mouse models of pulmonary hypertension. Br J Pharmacol, 2009. 157(7): p. 1241–9.

76. Peinado, V.I., et al., Expression of BK(Ca) channels in human pulmonary arteries: relationship with remodeling and hypoxic pulmonary vasoconstriction. Vascul Pharmacol, 2008. 49(4-6): p. 178–84.

77. Cogolludo, A., et al., Role of reactive oxygen species in Kv channel inhibition and vasoconstriction induced by TP receptor activation in rat pulmonary arteries. Ann N Y Acad Sci, 2006. 1091: p. 41–51.

