## supplemental figures for "Direct Effects of Cigarette Smoke in Pulmonary Arterial Cells alter Vascular Tone through Arterial Remodeling and Kv7.4 Channel Dysregulation"

**Supplementary table S1. List of the primer pairs used for gene expression analysis by real-time PCR.**

| Gene | 5′-3′ forward primer sequence | 5′-3′ reverse primer sequence |
|---|---|---|
| *ACTB* | GGCACCCAGCACAATGAAG | CCGATCCACACGGAGTACTTG |
| *CDKN1A* | CTGGAGACTCTCAGGGTCGAA | GCGGATTAGGGCTTCCTCTT |
| *CDKN2A* | GTGGACCTGGCTGAGGAG | CTTTCAATCGGGGATGTCTG |
| *IL6* | GACAGCCACTCACCTCTTCA | CCTCTTTGCTGCTTTCACAC |
| *CXCL8* | GCTCTGTGTGAAGGTGCAGT | CCAGACAGAGCTCTCTTCC |
| *NOX1* | TGCCCCTCAATCTCTCTCC | GGGACCATCCACTTCAATCC |
| *NOX4* | CCTCAACTGCAGCCTTATCC | CAACAATCTCCTGGTTCTCC |
| *CYBA* | TGGCGGGCGTGTTTGTGT | CCACGGCGGTCATGTACTTC |

**Supplementary figure S2. Contractile force generation is blocked in hPASMC after CSE treatment.**

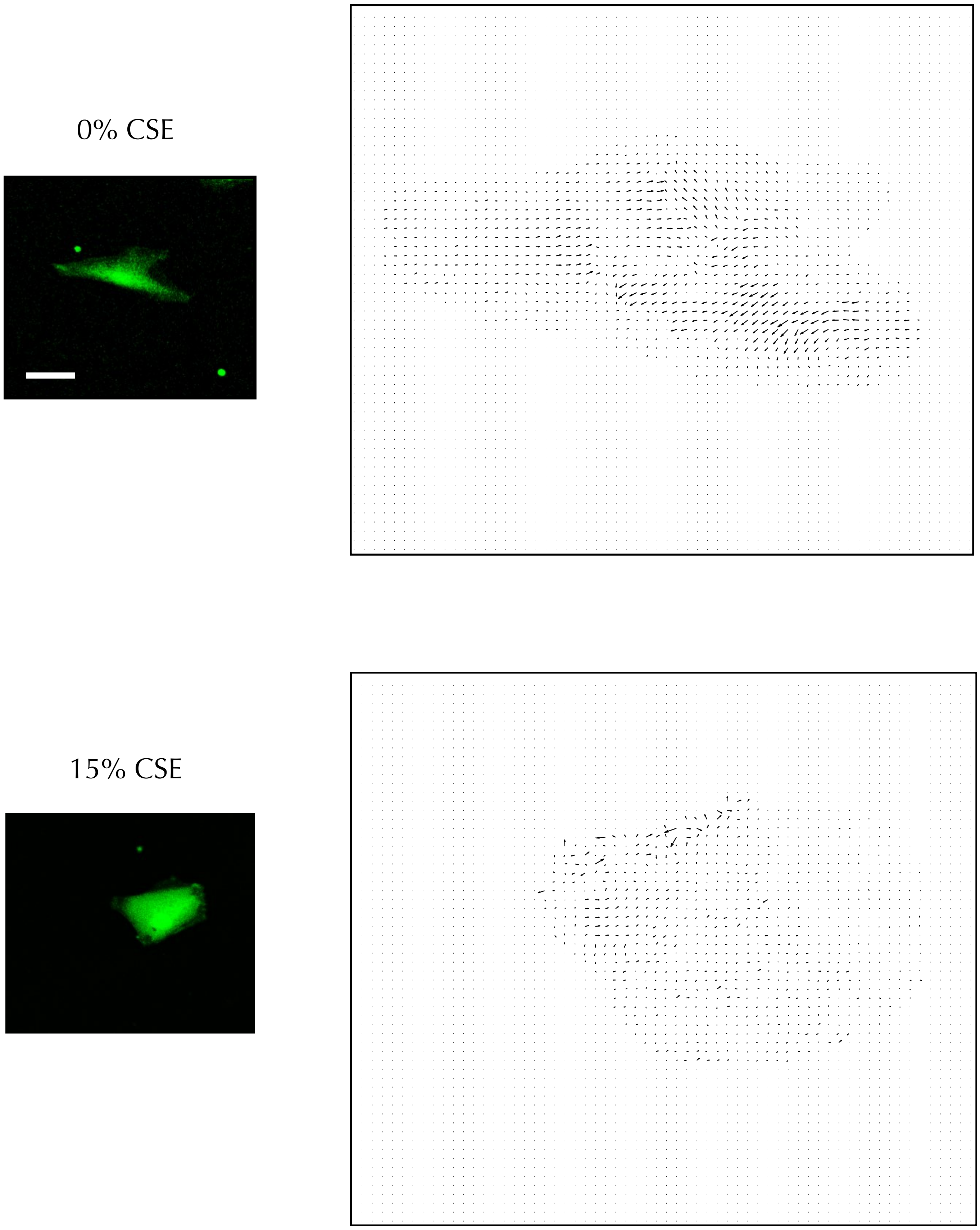

**Supplementary figure S3. Kv7.4 channel levels and activity are diminished in CSE-exposed pulmonary arteries.**

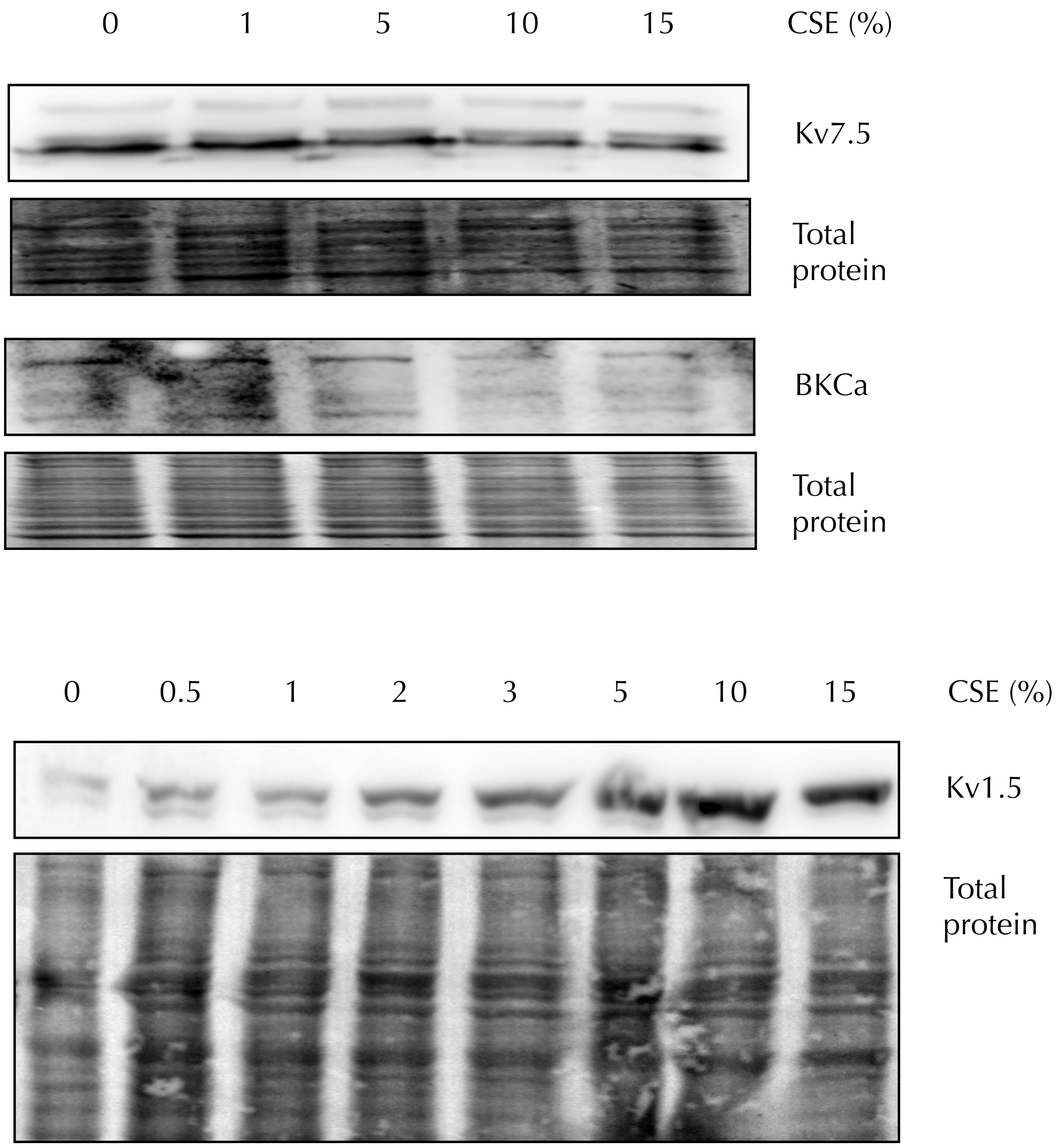
